# Batch culture and species effects modulate the decoupling between diatom frustule-bound and biomass nitrogen isotope signatures

**DOI:** 10.1101/2025.07.31.667865

**Authors:** Jochem Baan, Moritz F. Lehmann, Anja S. Studer

**Affiliations:** University of Basel, Department of Environmental Sciences, Bernoullistrasse 30, 4056 Basel, Switzerland

**Keywords:** biomass, diatom, frustule-bound organic compounds, isotope fractionation, nitrate, nitrogen stable isotopes

## Abstract

The stable nitrogen (N) isotope composition of organic matter encapsulated in diatom silica frustules (δ^15^N_DB_) from sedimentary records has been used as a proxy for reconstructing N consumption dynamics in the ocean over geologic timescales. This proxy relies on the assumption that δ^15^N_DB_ tracks biomass δ^15^N without being affected by internal N-isotope fractionation. However, recent ground-truthing efforts have shown that δ^15^N_DB_ can diverge from biomass δ^15^N values, though the extent and mechanisms behind this decoupling remain unclear.

In this study, we cultured two freshwater and two marine diatom species in batch cultures to test whether δ^15^N_DB_ (1) is subject to species-dependent internal ^15^N fractionation, and (2) reflects the δ^15^N of source nitrate to the same extent as biomass δ^15^N values, assessing potential asynchronous integration of the N isotope signal. We monitored the N-isotope systematics during diatom growth by measuring δ^15^N values of nitrate, bulk biomass and frustule-bound organic N throughout batch culture progression. We found that δ^15^N_DB_ did not follow typical Rayleigh fractionation dynamics and remained relatively stable, while biomass δ^15^N increased predictably with progressive fractional nitrate consumption. The observed divergence could only be partially explained by asynchronous integration of source-nitrate δ^15^N values into biomass versus frustule-bound organic N (i.e., delayed incorporation into frustule-bound material), as newly formed frustules predominantly recorded the δ^15^N of ^15^N-labeled nitrate added during growth. This demonstrates that δ^15^N_DB_ values capture the isotopic signature of newly assimilated nitrate rather than N derived from internal, or legacy, pools. We hypothesize that shifts in growth conditions during batch culture progression alter the coupling between carbon and nitrogen metabolism, leading to physiologically driven variation in internal ^15^N fractionation and corresponding offsets between δ^15^N_DB_ and biomass δ^15^N. Such sensitivity to internal isotope fractionation during biosynthesis implies that the interpretation of sedimentary δ^15^N_DB_ records is more intricate than previously assumed.

## 1. Introduction

The nitrogen (N) isotopic composition (δ^15^N) of diatom frustule-bound organic matter (δ^15^N_DB_) from sedimentary archives has been used as a promising proxy indicator for reconstructing nitrate utilization across various timescales in the ocean (Sigman et al., 1999; Robinson et al., 2004; Ren et al., 2015; Studer et al., 2015; Ai et al., 2020), and more recently, in lacustrine environments for paleoenvironmental reconstructions (Studer et al., 2024). The main advantage of this proxy is that δ^15^N_DB_ is thought to be physically protected from diagenetic alteration and bacterial degradation, unlike the δ^15^N of organic matter in bulk sediment which is known to be susceptible to post-depositional alterations (Lehmann et al., 2002; Robinson et al., 2012). Despite the fact that the δ^15^N_DB_ proxy has been applied in palaeoceanographic research for more than two decades, the mechanisms by which the δ^15^N signature of assimilated inorganic N is transferred into diatom biomass, and ultimately preserved in δ^15^N_DB_, remain poorly understood.

When diatoms grow, their cell walls become silicified, forming opaline frustule structures. This silicification occurs after mitosis, through the synthesis of new valves, and during cell expansion, when girdle bands are formed (Kröger and Poulsen, 2008). Frustule formation is facilitated by several N-containing organic compounds (e.g., silaffins, silacidins, and polyamines), which act as organic templates for silicate precipitation (Poulsen et al., 2003; Bridoux and Ingalls, 2010; Richthammer et al., 2011). These compounds become encapsulated within the frustule and can persist over long time scales. The N in these frustule-bound organic compounds originates from the intracellular organic N pool in the diatom, which in turn derives from the assimilation of inorganic N in the surrounding water. Under this framework, the δ^15^N values of assimilated inorganic N should be transferred into the cellular organic N pool and therefore reflected by δ^15^N_DB_. Accordingly, δ^15^N_DB_ values have been interpreted as proxies for nitrate utilization (Robinson et al., 2005; Brunelle et al., 2007), nitrogen cycling in general (Kalansky et al., 2011), and N eutrophication (Studer et al., 2024).

Efforts to ground-truth the diatom frustule-bound δ^15^N paleo-proxy through controlled batch culture experiments have shown that δ^15^N_DB_ values can differ substantially from biomass δ^15^N values (Horn et al., 2011; Jones et al., 2022; Robinson et al., 2025). Horn et al. (2011) found δ^15^N_DB_ values lower than those of biomass, with species-specific offsets, whereas Jones et al. (2022) reported the opposite pattern. More recently, Robinson et al. (2025) observed both positive and negative offsets, depending on species, and suggested that the nitrate-to-silicate uptake ratio may play a modulating role. Overall, these findings suggest that δ^15^N_DB_ values can vary substantially and do not consistently track biomass δ^15^N values, and that the underlying drivers of the offset remain poorly understood.

Culturing diatoms under controlled environmental conditions enables detailed observation of N isotope transfer from source nitrate into diatom biomass and frustule-bound material. In batch cultures, the δ^15^N values of nitrate and accumulated biomass typically follow Rayleigh-type substrate-product dynamics (Mariotti et al., 1981; Waser et al., 1998). Once assimilated, N is processed into various proteins and other N-containing biomolecules, including those found in diatom frustules (Sumper and Brunner, 2008; Kröger and Poulsen, 2008; Bridoux and Ingalls, 2010; Richthammer et al., 2011; Kirkham et al., 2017). As diatoms synthesize frustules over the course of a batch culture, the frustule-bound organic N pool accumulates. It might therefore be expected that δ^15^N_DB_ values reflect the integrated isotopic composition of the accumulating frustule-bound organic matter, similar to the integrated δ^15^N signal in bulk biomass (Mariotti et al., 1981). However, this expectation rests on two key assumptions: (1) that ^15^N fractionation associated with the synthesis of frustule-bound organic compounds is constant, and (2) that the integration time window of the isotope-signal incorporation from the inorganic N source is the same for biomass and frustule-bound organic N. With regards to the first assumption, internal N-isotope fractionation during biosynthesis may vary due to metabolism-driven biochemical reactions (Werner and Schmidt, 2002; Tcherkez and Hodges, 2008). Additionally, the δ^15^N values of amino acids (the building blocks of proteins and polyamines) can vary between compounds and across species (Macko et al., 1987; Werner and Schmidt, 2002), suggesting that δ^15^N_DB_ values may also depend on (species-specific) differences in the frustule-bound organic N matrix/composition (Sumper and Brunner, 2008; Bridoux and Ingalls, 2010). As for the second assumption, frustule-bound organic N may derive from an internal N pool with a distinct isotopic signature, potentially leading to asynchronous δ^15^N-signal integration relative to bulk biomass. To date, neither of these assumptions have been experimentally verified, yet the observation that δ^15^N_DB_ values can be both higher and lower than biomass δ^15^N values (Horn et al., 2011; Jones et al., 2022; Robinson et al., 2025) implies that at least one of them may not hold. As such, it remains uncertain whether δ^15^N_DB_ values consistently reflect biomass and/or nitrate δ^15^N values, and thus exhibit predictable Rayleigh-type fractionation behavior.

Diatom culture studies aimed at ground-truthing the δ^15^N_DB_ proxy have so far exclusively focused on marine species. However, since the proxy has recently also been applied in lacustrine systems (Studer et al., 2024), it is important to investigate the relationship between nitrate, biomass and frustule-bound δ^15^N values in freshwater diatoms. Moreover, freshwater species generally exhibit higher silicate uptake rates compared to most marine species (Conley et al., 1989). Therefore, further insight into whether and how silicate-to-nitrate uptake ratios influence δ^15^N_DB_ relative to biomass δ^15^N (Robinson et al., 2025) may be gained by including freshwater taxa in the experimental design.

In this study, we cultured two marine and two freshwater diatom species under controlled environmental conditions, with nitrate as the limiting nutrient. These batch cultures were closely monitored, and δ^15^N values of nitrate, bulk biomass and frustule-bound organic N were monitored throughout the progression of each culture. This experimental setup allowed us to specifically test whether variation in frustule-bound δ^15^N could be explained by that of nitrate and/or biomass, and whether the observed patterns were consistent (1) among species, (2) between ecosystems (freshwater vs. marine), and (3) over time during batch culture progression. In a separate ^15^N-labeling experiment we manipulated the δ^15^N of the nitrate source during growth, to assess if bulk-biomass and frustule-bound δ^15^N values capture the nitrate isotope signal concurrently, or whether differences in isotopic integration timing exist between these N pools. Overall, our results highlight that δ^15^N_DB_ values do not strictly follow Rayleigh-type dynamics in batch culture. However, this deviation from expected patterns is not due to (species-specific) differences in the timing of nitrate-δ^15^N incorporation into diatom biomass versus frustule-bound N. Instead, our findings suggest that the observed offset is more likely driven by variable, growth-condition-dependent biochemical ^15^N fractionation.

## 2. Methods

### 2.1. Diatom cultures

Two colony-forming freshwater species were selected: the centric *Aulacoseira granulata* and the pennate *Fragilaria capucina*, both isolated from Swiss lakes and kindly provided by Marta Reyes (Swiss Federal Institute for Aquatic Research, Dubendorf, Switzerland). Two marine diatom species forming individual cells were obtained from the Culture Collection of Algae & Protozoa (CCAP, Scottish Association for Marine Science, Dunbeg, Scotland, United Kingdom): *Chaetoceros calcitrans* (CCAP 1085/3), a small centric diatom, and *Coscinodiscus radiatus* (CCAP 1013/11), a large centric diatom.

The growth medium for freshwater diatoms was based on the CCAP ‘Diatom Medium’ recipe, with some modifications, resulting in the following starting concentrations: 102 µM Ca(NO_3_)_2_, 100 µM K_2_HPO_4_, 101 µM MgSO_4_, 200 µM KHCO_3_, 200-350 µM Na_2_SiO_3_, 36 µM Fe-EDTA, 5 µM MnCl_2_, 0.2 µM CuSO_4_, 0.9 µM ZnSO_4_, 0.5 µM Na_2_MoO_4_, 40 µM H_3_BO_3_, and vitamins (0.04 mg/L cyanocobalamin, thiamine HCl, and biotin each).

The medium for marine diatoms was based on the ESAW artificial seawater recipe (Harrison et al., 1980; Berges et al., 2001), with some modifications. The base seawater composition followed ESAW with the exception that NaBr was used instead of KBr. Micronutrient composition also followed ESAW, except CoSO_4_ was replaced with CoCl. Since copper is not typically included in ESAW, but was added to the freshwater medium, 0.1 µM of CuSO_4_ was also added here. Iron was provided as 36 µM Fe-EDTA. The same vitamin concentrations were used as in the freshwater medium. Lastly, nitrate, phosphate and silicate were added at starting concentrations of 254 µM Ca(NO_3_)_2_, 103 µM K_2_HPO_4_ and 150-350 µM Na_2_SiO_3_. In order to obtain sufficient frustule material from *C. calcitrans*, which is weakly silicifying, a higher initial nitrate concentration (∼508 µM) was used. The pH of the artificial seawater medium was adjusted to 8.1 ± 0.1 using a 0.5 M NaOH prior to filtration.

All media were filter-sterilized using sterile 0.2 µm polyether sulfone (PES) membranes into autoclaved containers. Since the Na_2_SiO_3_ stock solution had a high pH (12-13), its addition influenced the pH of the medium and caused precipitation and filtration problems. Therefore, silicate was added aseptically *after* filtration. In some freshwater batch cultures, 400 mg/L N-Tris(hydroxymethyl)methyl-2-aminoethanesulphonic acid (TES) buffer was added to stabilize pH during growth, and to counteract the pH rise from Na_2_SiO_3_ addition (Table 1). For marine cultures and freshwater batch cultures 5 & 6, silicate was added as a neutralized colloid (Table 1): a 50 mM Na_2_SiO_3_ stock solution was titrated with 1M HCl to pH 7.5-8.5 while stirring, forming a Na_2_SiO_3_ colloid that was then aseptically added post-filtration.

**Table 1:** Overview of initial and final nitrate concentrations, Si:NO_3_^-^ uptake ratios, frustule N contents, C and N contents, and batch culture durations for experiments with different diatom cultures. Values are reported as mean ± SD where applicable. bSi, biogenic silica

| Species | No. | Starting<br>[NO<sub>3</sub><sup>-</sup>]<br>(uM) | Final<br>[NO<sub>3</sub><sup>-</sup>]<br>(uM) | Si:NO<sub>3</sub><sup>-</sup><br>uptake<br>ratio | Frustule N<br>content<br>(nmol N mg<sup>-1</sup><br>bSi) | Biomass C<br>fraction (g g<sup>-1</sup>) | Biomass N<br>fraction (g g<sup>-1</sup>) | Number<br>of<br>frustule<br>samples | Culture<br>duration<br>(d) | Notes |
| --- | --- | --- | --- | --- | --- | --- | --- | --- | --- | --- |
| *Aulacoseira granulata* | 1 | 195 | 56 | 2.6 | 14.2 ± 1.6 | 0.19 ± 0.02 | 0.031 ± 0.001 | 3 | 13 | |
| | 2 | 196 | 14 | 2.1 | 12.4 ± 0.5 | 0.22 ± 0.04 | 0.035 ± 0.002 | 3 | 13 | |
| | 3 | 191 | 0 | 2.6 | 12.8 ± 0.7 | 0.20 ± 0.02 | 0.037 ± 0.006 | 4 | 14 | With TES buffer |
| | 4 | 196 | 0 | 1.8 | 15.7 ± 5.9 | 0.18 ± 0.02 | 0.035 ± 0.004 | 5 | 15 | With TES buffer |
| | 5 | 203 | 0 | 2.3 | 9.0 ± 0.5 | 0.16 ± 0.01 | 0.031 ± 0.003 | 6 | 15 | <sup>15</sup>N-labeled nitrate addition |
| | 6 | 208 | 1 | 1.9 | 9.9 ± 0.7 | 0.18 ± 0.02 | 0.033 ± 0.002 | 6 | 12 | <sup>15</sup>N-labeled nitrate addition |
| *Fragilaria capucina* | 1 | 194 | 45 | 2.2 | 13.2 ± 1.0 | 0.23 ± 0.01 | 0.030 ± 0.002 | 3 | 22 | |
| | 2 | 194 | 21 | 2.3 | 14.3 ± 2.6 | 0.26 ± 0.03 | 0.032 ± 0.001 | 3 | 21 | |
| | 3 | 185 | 1 | 2.9 | 9.3 ± 1.3 | 0.20 ± 0.03 | 0.028 ± 0.002 | 5 | 15 | With TES buffer |
| | 4 | 194 | 6 | 2.6 | 8.8 ± 1.2 | 0.19 ± 0.03 | 0.028 ± 0.002 | 6 | 17 | With TES buffer |
| | 5 | 205 | 2 | 1.5 | 7.1 ± 2.3 | 0.21 ± 0.03 | 0.030 ± 0.005 | 6 | 21 | <sup>15</sup>N-labeled nitrate addition |
| | 6 | 200 | 0 | 1.9 | 6.0 ± 1.3 | 0.22 ± 0.02 | 0.028 ± 0.003 | 6 | 21 | <sup>15</sup>N-labeled nitrate addition |
| *Chaetoceros calcitrans* | 1 | 512 | 4 | 0.5 | 17.6 ± 1.5 | 0.15 ± 0.03 | 0.028 ± 0.005 | 4 | 8 | |
| | 2 | 502 | 4 | 0.4 | 15.6 ± N/A | 0.14 ± 0.06 | 0.025 ± 0.011 | 4 | 8 | |
| | 3 | 500 | 2 | 0.7 | 4.1 ± 0.7 | 0.13 ± 0.04 | 0.027 ± 0.007 | 4 | 7 | |
| *Coscinodiscus radiatus* | 1 | 509 | 27 | 4.1 | 2.2 ± 0.4 | 0.11 ± 0.01 | 0.016 ± 0.002 | 5 | 47 | |
| | 2 | 513 | 2 | 4.4 | 1.7 ± 0.3 | 0.12 ± 0.02 | 0.016 ± 0.002 | 5 | 47 | |

Diatom culture stocks were maintained in 100-200 mL Erlenmeyer flasks, filled with 50-100 mL of medium and covered with aluminum foil. Cultures were grown at 20 °C under a 12h/12h light/dark cycle with ∼140 µmol m^−2^ s^−1^ photons from LED lighting in the growth chambers (MLR-350, SANYO, Japan). Stocks were manually stirred daily, and refreshed every 2-4 weeks, either by addition of medium, or by transferring 5 mL of culture into fresh 50 mL medium in a new flask.

The batch culture experiments were conducted in 20-L clear polycarbonate carboys filled with sterile-filtered medium. Each carboy was fitted with three ports: one for sampling, one for air inflow to aerate the medium, and one for air outflow. Air was supplied via a compressed air system through 0.2 µm hydrophobic PTFE filters. Carboys were placed in a large custom-built walk-in phytotron growth chamber at 20 °C, with ceiling-mounted LEDs providing ∼140 µmol m^−2^ s^−1^ photons. A 12h/12h light/dark cycle was used, with gradual light-intensity ramp-up and ramp-down over the first and last hour of the light period, respectively. Carboys were equilibrated with ambient air for at least 24 h prior to inoculation. Cultures were inoculated with 20-40 mL of recently transferred stock culture. This relatively small inoculum delayed biomass sample collection in early incubation stages but minimized disruption of expected closed-system isotope dynamics. Each species was cultured in at least two replicate batch cultures.

Media were regularly sampled to monitor growth, pH, and nutrient (i.e., nitrate and silicate) consumption. Growth was tracked via *in vivo* fluorescence using an AquaFluor handheld fluorometer (Turner Designs, CA, USA). Samples were filtered through 0.22 µm PES syringe filters. In freshwater cultures, nitrate concentrations were measured by spectrophotometric analysis using LCK339 nitrate test kits (Hach, Germany). In artificial seawater, nitrate was measured using a chemiluminescence method (described below) due to interference from high ion concentrations with the nitrate test. For dissolved silica analysis, filtered samples were acidified to ∼pH 2 using 1 µL 6N HCl per mL of sample. Silicate concentrations were determined photometrically (Strickland and Parsons, 1972), using molybdate-3, citric acid, and amino acid F reagents, and calibrated against a 1 mg/l SiO_2_ standard (Hach, Germany). When silicate became limiting before a substantial nitrate fraction was consumed, additional silicate was added to the batch culture. Samples for nitrate concentration and δ^15^N analysis were stored frozen at -20 °C. Silicate samples were stored at room temperature to avoid polymerization at high silicate concentrations.

Once biomass reached sufficient density – based on nitrate and silicate consumption – 1-4 L samples were harvested at regular time intervals. Biomass was concentrated by centrifugation and split into two aliquots: one for bulk biomass δ^15^N and δ^13^C measurements and elemental (N and C) analysis, and one for the analysis of frustule-bound δ^15^N values. Bulk-biomass aliquots were frozen at -20 °C, freeze-dried, and stored in the dark at room temperature prior to analysis. Frustule-bound δ^15^N samples were stored frozen at -20 °C until further processing.

Two replicate batch-cultures of each freshwater diatom species (culture numbers 5 & 6; Table 1) were grown as described above, except that after the first 2-4 biomass samples had been harvested, and once nitrate had decreased to ∼50 % of its initial concentration, a 4 mM solution of ^15^N-labelled nitrate (δ^15^N ≈ 4000 ‰) was added. The volume added was calculated based on mass-balance targeting an 80-100 ‰ increase in nitrate δ^15^N, with minimal change in nitrate concentration (1-2 μM). After the ^15^N-nitrate spike, cultures were monitored and harvested regularly until all nitrate was consumed. In this experiment, the relative abundance of empty frustules in harvested biomass was assessed microscopically to determine whether a potential decoupling between biomass and frustule-bound δ^15^N value could result from an increased contribution of older, unlabeled frustules.

### 2.2. Sample processing and isotope analysis

#### 2.2.1. Nitrate concentration measurements

Nitrate concentrations were determined using a chemiluminescence method. Briefly, samples were injected into a heated 0.1 M vanadium(III) in 2N HCl solution to convert sample nitrate to NO (Braman and Hendrix, 1989), followed by chemiluminescent detection of the NO gas after reaction with ozone in a TELEDYNE T200 NO/NO_2_/NOx analyzer (TELEDYNE API, CA, USA). Sample nitrate concentrations were determined by comparison to multiple injections of KNO_3_ reference standards of known nitrate concentrations.

#### 2.2.2. Biomass δ^15^N and δ^13^C values, and N and C content

Aliquots of 1-3 mg of freeze-dried diatom biomass were weighed into tin cups. Biomass δ^15^N and δ^13^C values, as well as total N and C contents were determined using a Delta V Plus isotope ratio mass spectrometer (IRMS; Thermo Fisher Scientific) coupled to a Flash Elemental Analyzer (EA). Scale normalization for δ^15^N values was performed using USGS61 (δ^15^N = -2.87 ‰; δ^13^C = -35.05 ‰), and USGS62 (δ^15^N = 20.17 ‰; δ^13^C = -14.79 ‰) international reference materials. For samples from the ^15^N-labeled nitrate addition experiments, we additionally included USGS41 (δ^15^N = 47.57 ‰; δ^13^C = 37.63 ‰). Scale normalization for δ^13^C values was based on USGS40 (δ^15^N = -4.52 ‰; δ^13^C = -26.39 ‰), USGS62, and USGS64 (δ^15^N = 1.76 ‰; δ^13^C = - 40.81 ‰) standards. Elemental N and C contents were determined from IRMS peak areas using a calibration curve that was established based on the analysis of a wheat flour standard (1.36 % N; 39.38 % C; Säntis analytical, Switzerland) and EDTA (9.58 % N; 41.06 % C). Analytical reproducibility was assessed by repeated analysis (n = 5) of biomass from a commercial diatom culture (*Thalassiosira weissflogii*, TW1800, Reed Mariculture, CA, USA), yielding mean ± SD δ^15^N and δ^13^C values of 6.9 ± 0.3 ‰ and -10.50 ± 0.1 ‰, respectively.

#### 2.2.3. Frustule-bound and ambient nitrate δ^15^N analysis

All glassware used in frustule-bound δ^15^N analysis was cleaned by combustion in a muffle oven at 450 °C for 6 h. Diatom frustules were cleaned as described by (Morales et al., 2013). Briefly, diatom samples underwent at least two cycles of sonication in 2 % (w/v) sodium dodecyl sulfate (SDS), followed by centrifugation, decanting, sonication in purified water, centrifugation and decanting. Non-frustule-bound biomass was then further removed through subsequent steps of oxidation by sulfuric acid and potassium permanganate (KMnO_4_), 14 % perchloric acid for 1 h at 100 °C, and 70 % perchloric acid for 1 h at 100 °C. Cleaned diatom frustules were collected on 2 µm polycarbonate filter membranes and rinsed thoroughly with purified water. Diatom samples were then dried at 60 °C in a dedicated, low-N oven. Dissolution of the opal frustules and oxidation of the released frustule-bound organic N was performed within one week of drying.

Approximately 2-7 mg of dried frustules were weighed into 4 mL glass vials, but samples with lower weights were transferred to glass vials using purified water. Persulfate oxidizing reagent (POR) solution used to dissolve diatom frustules and oxidize organic N to nitrate was added (1.5ml per vial). POR consisted of potassium persulfate (5% w/v) and sodium hydroxide (5% w/v). The potassium persulfate salt was pre-purified to remove residual N via multiple dissolution and crystallization cycles.

Vials were autoclaved for 90 minutes at 121 °C. Following oxidation, sample pH was adjusted to 4 - 7 using ultrapure HCl. Nitrate concentrations of oxidized frustule samples were determined using the chemiluminescence method described above and the δ^15^N of the frustule-derived nitrate was measured using the denitrifier method (see below; Sigman *et al*., 2001). Each sample batch included vials with known concentrations of amino acid standards (USGS64, USGS65, USGS40 and USGS41) and procedural blanks to assess recovery, oxidation completeness, and blank contribution. Procedural blank size and δ^15^N values were used for post-correction of frustule-bound δ^15^N values using isotope mass-balance considerations.

Consistency of the frustule-cleaning and oxidation method was assessed by routinely including material from a commercial diatom culture of *T. weissflogii* (TW1800, Reed Mariculture). This regularly co-processed material yielded δ^15^N_DB_ values of 0.4 ± 0.3 ‰ (n = 10). Additionally, to further determine the consistency of the oxidation method, diatom frustule material from two reference sources - a commercial *Thalassiosira pseudonana* culture (TP1800, Reed Mariculture) and a phytoplankton sample collected in Lake Murten, Switzerland (2023) - was cleaned once in bulk, stored in sealed vials isolated from the atmosphere, and regularly co-analyzed across runs. These samples yielded frustule-bound δ^15^N values of -6.1 ± 0.5 ‰ (n = 9), and 8.4 ± 0.4 ‰ (n = 8), respectively. Across all diatom culture samples, the mean procedural blank contribution was 1.5 %, with a maximum of 5.6 %. Corresponding corrections to frustule-bound δ^15^N values were, on average, 0.13 ‰ with a maximum of 0.63 ‰.

Ten nanomoles of dissolved nitrate (from culture medium or from oxidized diatom frustules) was converted to N_2_O using the denitrifier method (Sigman et al., 2001). The δ^15^N of the resulting N_2_O was measured by gas-chromatography isotope ratio mass spectrometry (GC-IRMS; Delta V Plus, Thermo Fisher Scientific) coupled to a Gasbench used for N_2_O purification. Scale normalization was performed using nitrate reference materials with known δ^15^N: USGS34 (−1.8 ‰), IAEA N3 (4.7 ‰) and a lab-internal standard UBN-1 (14.15 ‰). For the analysis of nitrate samples from the ^15^N-labelling experiment, the USGS32 standard (δ^15^N = 180 ‰) was also included. Analytical precision was assessed through repeated analyses of nitrate from a Deep-Pacific-Ocean water sample (5.2 ± 0.1 ‰, n = 28).

### 2.3. Data analysis

All δ^15^N values are reported relative to AIR-N_2_, and δ^13^C values relative to the Vienna Pee Dee Belemnite (V-PDB) standard. All calculations and statistical analyses were performed in R version 4.3.0 (R Core Team, 2023). For each batch culture, the nitrate-to-silicate uptake ratio was determined from the slope of a linear regression between nitrate and silicate concentrations, using only data points collected before the addition of supplemental silicate to the culture.

Assuming that ^15^N fractionation during nitrate assimilation in the closed-system batch cultures follows Rayleigh-type behavior/systematics, the N-isotope fractionation (i.e., isotope effect), ε_biomass/NO3_, was estimated as the slope of the linear relationship between the natural logarithm of the N isotopic ratio of nitrate (^15^N/^14^N) and the fraction of nitrate remaining in the medium (*f*) (Mariotti et al., 1981). This relationship was modelled with ordinary least squares regression (OLS) using the *lm* function in R. The intercept of this regression approximates the initial nitrate δ^15^N value, i.e., when *f* = 1 (δ^15^N*_f_*_1_). Using ε_biomass/NO3_ and δ^15^N*_f_*_1_ we calculated the expected δ^15^N value of biomass (i.e., the accumulated product) as a function of *f*, using the Rayleigh-based equation (Mariotti et al., 1981):

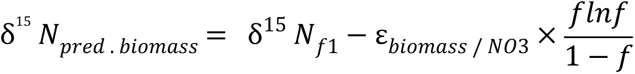

To evaluate whether the integration of nitrate δ^15^N values into biomass followed Rayleigh dynamics predictably, we tested if the slope of the relationship between predicted and measured biomass δ^15^N values differed significantly from 0 and 1. Additionally, to assess how faithfully biomass δ^15^N values are recorded in frustule-bound N, we tested whether the slope of the regression between δ^15^N values of frustule-bound N and those of bulk biomass differed significantly from 0 and 1. These (linear) regressions were also conducted using the *lm* function in R. The N-isotopic offset between δ^15^N values of frustule-bound N and biomass was calculated as ε_DB/biomass_ = α – 1, and α = (δ^15^N_DB_ + 1)/(δ^15^N_biomass_ + 1), expressed in ‰ (Coplen, 2011). We then related ε_DB/biomass_ values to the nitrate drawdown (*f*) to determine whether ε_DB/biomass_ changed with batch culture progression. Lastly, the slopes of these relationships were compared to ε_biomass/NO3_, but this analysis was limited to batch cultures with at least four ε_DB/biomass_ values (Table 1).

All the above analyses excluded samples from the ^15^N-labeled nitrate addition experiments. For these experiments, the biomass δ^15^N values after label addition were estimated by isotope mass balance, considering *f* values and the δ^15^N values of both the newly synthesized biomass (after addition of the ^15^N-labeled nitrate), and the pre-existing accumulated biomass (before ^15^N-label addition).

## 3. Results

### 3.1. Nutrient concentrations during batch cultures

Batch cultures lasted 12-15 days for *A. granulata*, 15-22 days for *F. capucina*, 7-8 days for *C. calcitrans*, and 47 days for *C. radiatus* (Table 1; Fig. S1 and S2). In 14 out of 17 batch-culture experiments, nitrate was consumed to concentrations corresponding to *f* values <0.1. Silicate-to-nitrate uptake ratios were highly variable among species: for the freshwater species they ranged from 1.5 and 2.9, for *C. calcitrans* from 0.4 and 0.7. For *C. radiatus* the highest Si:NO_3_ ^-^ uptake ratios were observed, ranging between 4.1 and 4.4 (Table 1).

### 3.2. Fractionation of nitrate N isotopes during nitrate assimilation

The isotope effect associated with nitrate assimilation (ε_biomass/NO3_), varied substantially among species (Fig. 1). The largest ε_biomass/NO3_ values were found for *C. radiatus* (−13.8 ±0.2 ‰), followed by *A. granulata* (−10.8 ±1.0 ‰), and *C. calcitrans* (−3.1 ±0.5 ‰), with the smallest values observed for *F. capucina* (−1.9 ±0.2 ‰).

**Figure 1:**
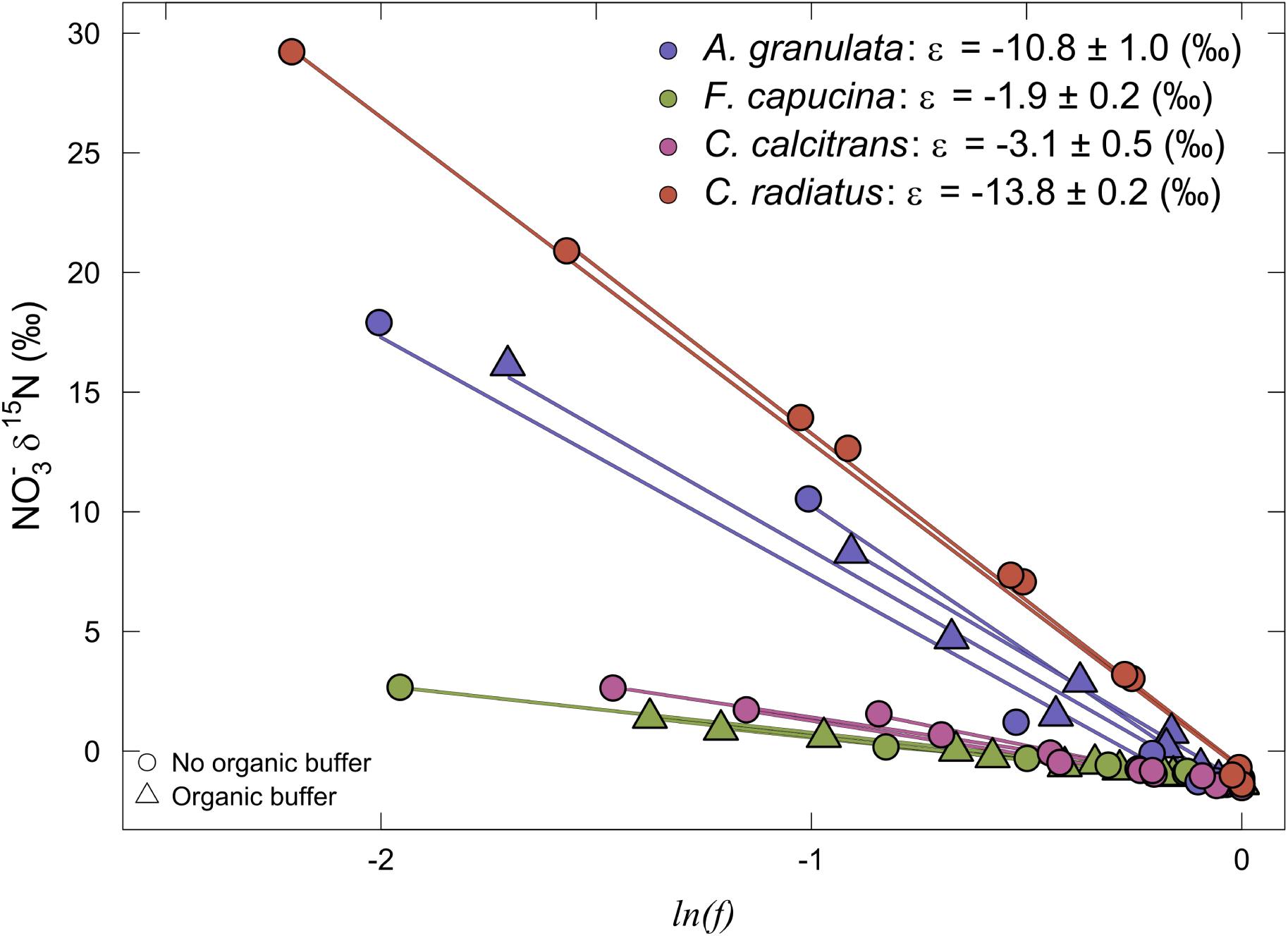
Nitrate ^15^N fractionation (ε_biomass/NO3_) during assimilation by different diatom species. ε values given in the legend represent species means ± SD, based on slopes of the linear regression, where the SD is derived from ε values obtained from culture replicates. Note that ε values were calculated from nitrogen isotope ratios rather than δ^15^N values; however, for ease of interpretation, the regressions are shown on the δ^15^N scale.

### 3.3. Biomass δ^15^N and δ^13^C values

Predicted biomass δ^15^N values – calculated using the cumulative-product equation by (Mariotti et al., 1981) and based on the observed nitrate isotope fractionation during nitrate assimilation – were very similar to measured biomass δ^15^N values (Fig. 2) with a regression slope of 0.95 (p < 0.001 for slope ≠ 0), close to but significantly different from 1 (p = 0.026 for slope ≠ 1). The explanatory power of this relationship was very high (R^2^ = 0.97), indicating that predicted biomass δ^15^N values explain nearly all of the observed variation in measured biomass δ^15^N values.

**Figure 2:**
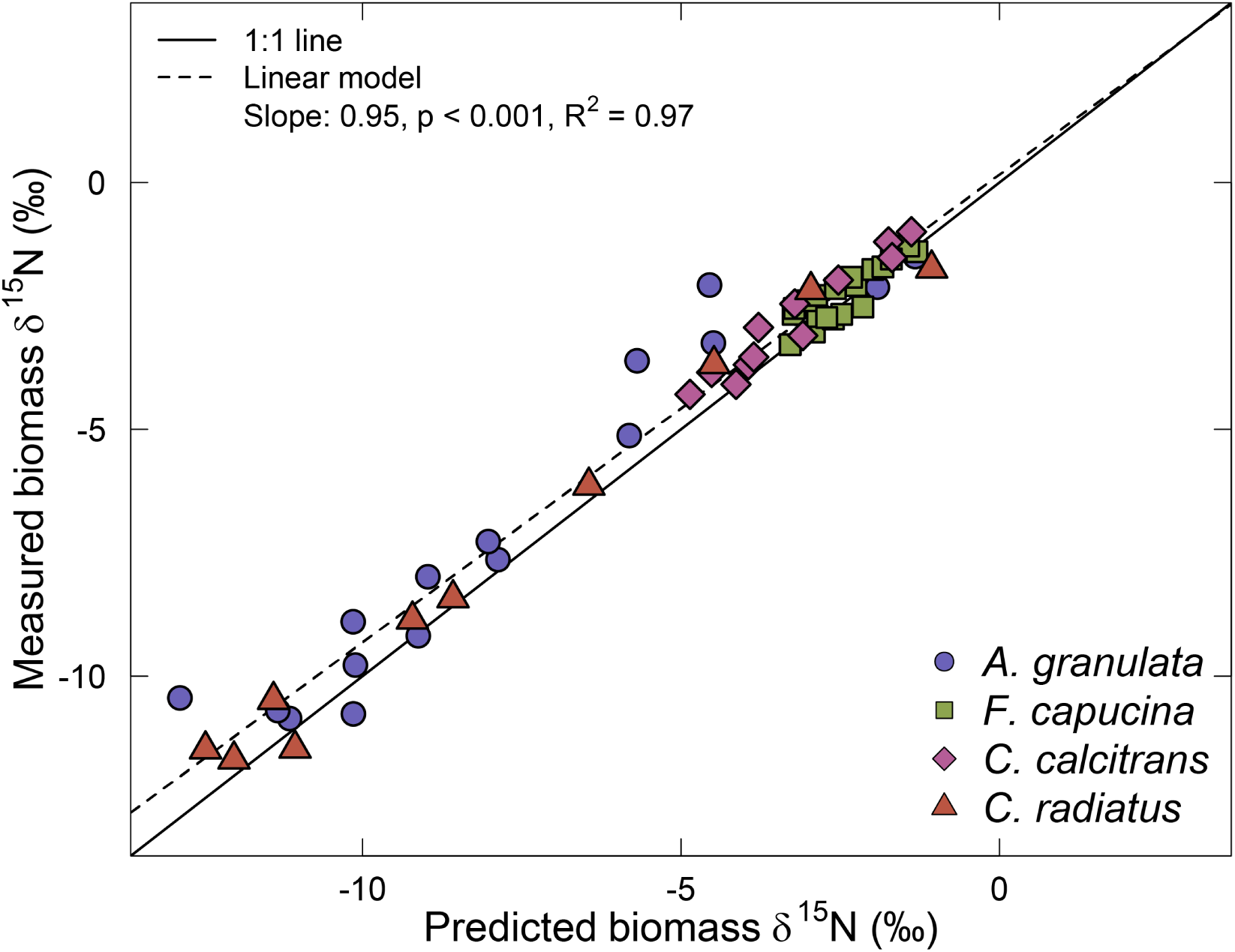
Comparison of predicted versus measured biomass δ^15^N values. Predicted biomass δ^15^N values were calculated using the cumulative-product equation (Mariotti et al. 1981), based on the observed ^15^N fractionation during nitrate assimilation and the fraction of nitrate remaining at the time of sampling.

Consistent with the species-specific ε_biomass/NO3_, biomass δ^15^N values at high *f* values were lowest in *A. granulata* and *C. radiatus*, around -11 to -12 ‰, while those of *F. capucina* and *C. calcitrans* were never lower than -5 ‰ (Fig. 3). Upon (near) complete consumption of nitrate in the medium, for all species, biomass δ^15^N approached values close to the starting nitrate δ^15^N value of -1.4 ‰ (Fig. 3).

**Figure 3:**
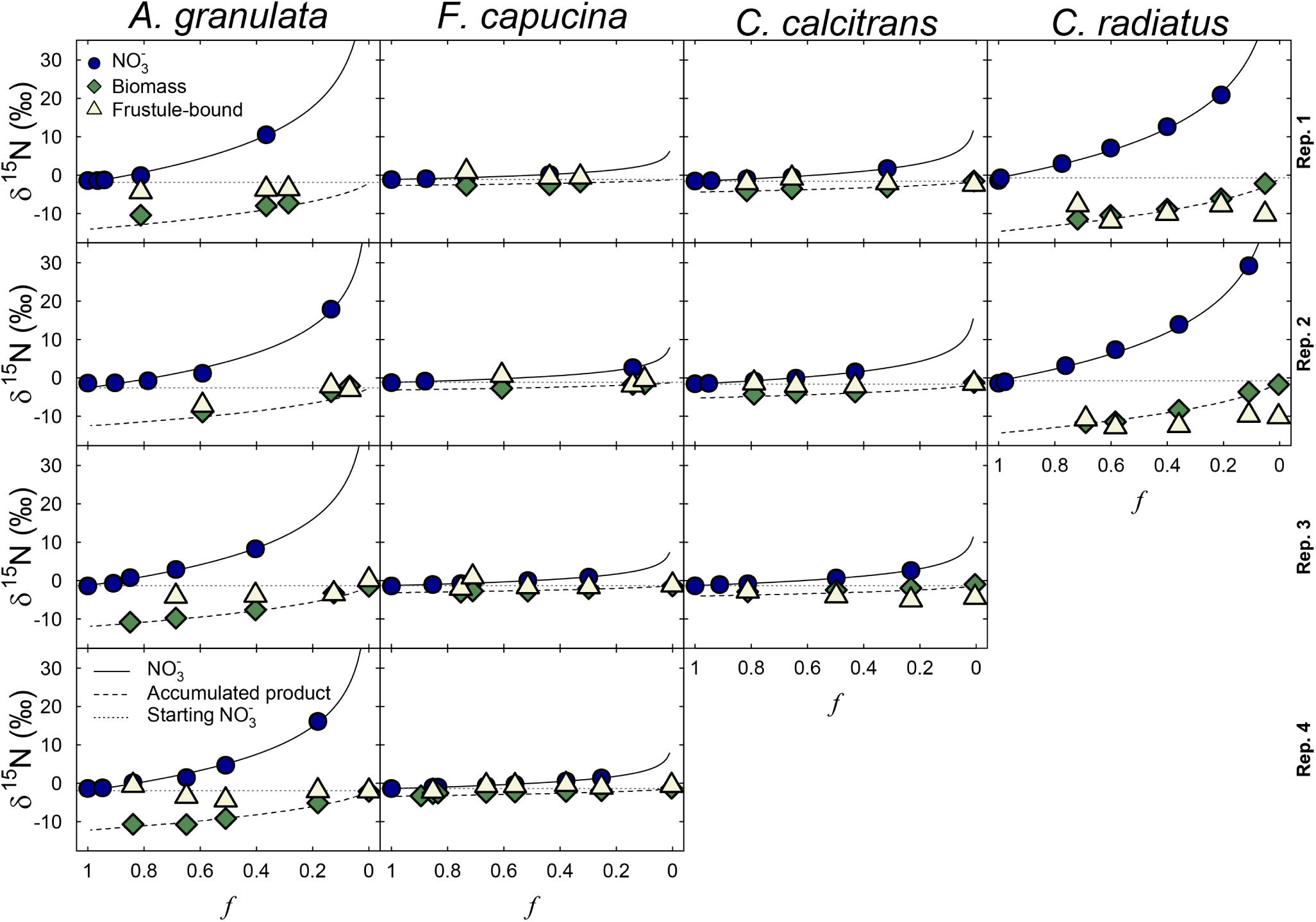
δ^15^N values of nitrate (blue circles), biomass (green diamonds) and frustule-bound N (white triangles) plotted against the fraction of nitrate remaining in the medium (f). Panels represent results from single batch cultures, organized by species (columns) and culture replicates (rows). Solid, dashed, and dotted lines represent theoretical δ^15^N values for nitrate, accumulated biomass, and starting nitrate, respectively. For A. granulata and F. capucina, culture replicates 3 and 4 were grown in medium containing TES buffer.

For *A. granulata*, *F. capucina* and *C. calcitrans*, biomass δ^13^C values increased with decreasing *f* (Fig. S3), with the magnitude of the increase being larger for species that more rapidly consumed the nitrate in the medium (Table 1; Fig. S1). For *C. radiatus*, there was no strong increase in biomass δ^13^C values during the batch culture, although the values were elevated compared to the lowest biomass δ^13^C values in the other species.

### 3.4. Diatom frustule-bound δ^15^N values

Linear regression of δ^15^N_DB_ values against biomass δ^15^N values across all species shows a significant relationship (p = 0.003 for slope ≠ 0; Fig. 4). However, the slope of this relationship was significantly different from 1 (p < 0.001 for slope ≠ 1), and the explanatory power was relatively low (R^2^ = 0.32). Moreover, both the significance and explanatory power of this relationship decreased when the data were analyzed at the species- or culture-specific level. This is further illustrated by increasing biomass δ^15^N values with decreasing *f*, while δ^15^N_DB_ values did not show substantial change (Fig. 3).

**Figure 4:**
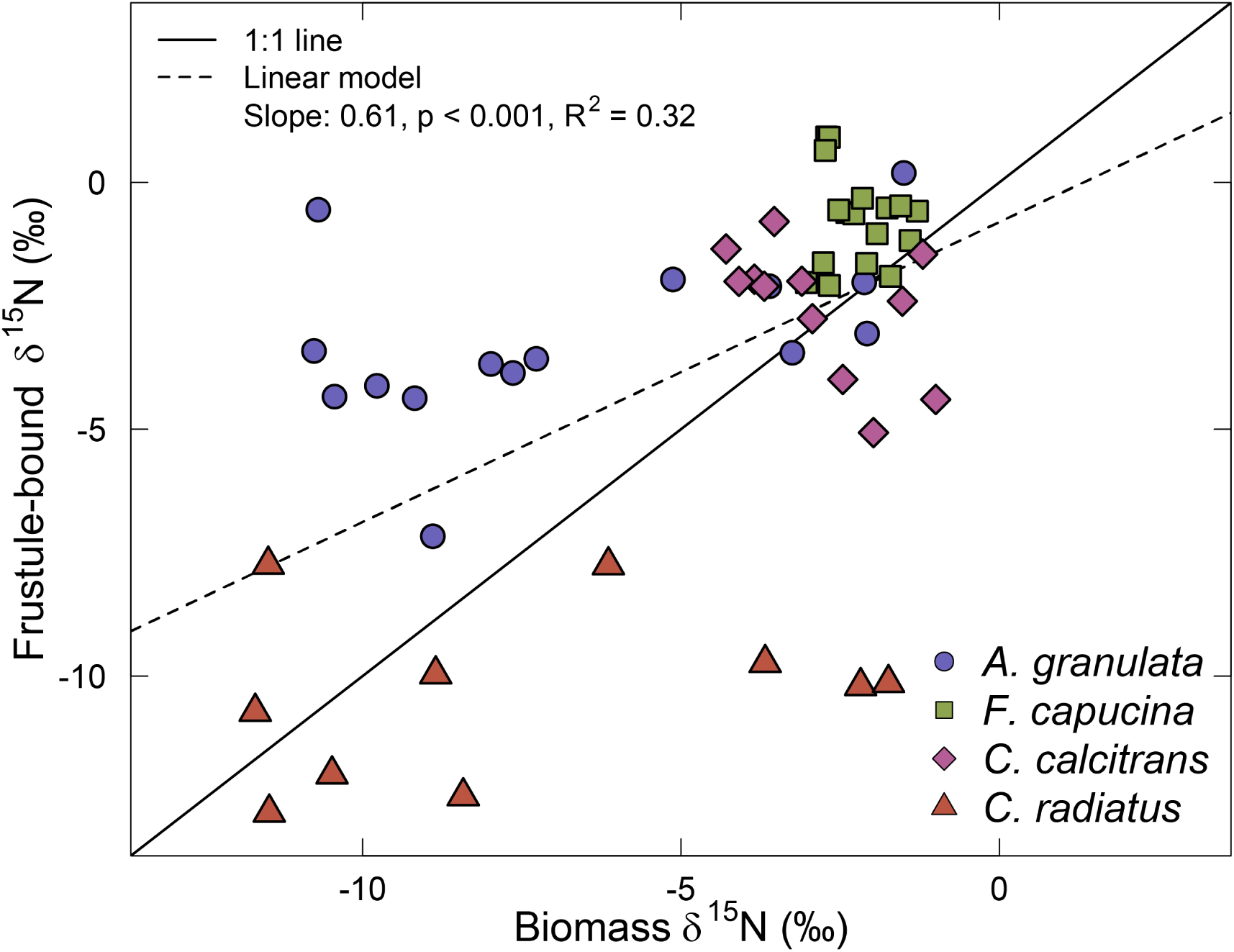
Comparison between frustule-bound and biomass δ^15^N values. The linear regression (dashed line) was determined across all species and batch cultures.

The isotopic offset between δ^15^N_DB_ values and biomass δ^15^N values, expressed as ε_DB/biomass_, was highly variable, from +10.3 to -8.4 ‰ depending on the species (Fig. 5). In nearly all cultures, ε_DB/biomass_ decreased with decreasing *f*. The magnitude of this decrease differed among species, with the strongest shifts observed for *A. granulata* and *C. radiatus*, and weaker decreases in *F. capucina* and *C. calcitrans*. In *A. granulata*, ε_DB/biomass_ decreased from +10 to +6 ‰ at *f* ≈ 0.8 to around 0 ‰ when *f* approached zero, whereas in *C. radiatus* it decreased from +5 to +1 ‰ at *f* = 0.7 to -8 ‰ when *f* approached zero. In *F. capucina* and *C. calcitrans*, ε_DB/biomass_ decreased by roughly 4 ‰ with *f* decreasing from 0.8 to 0.

**Figure 5:**
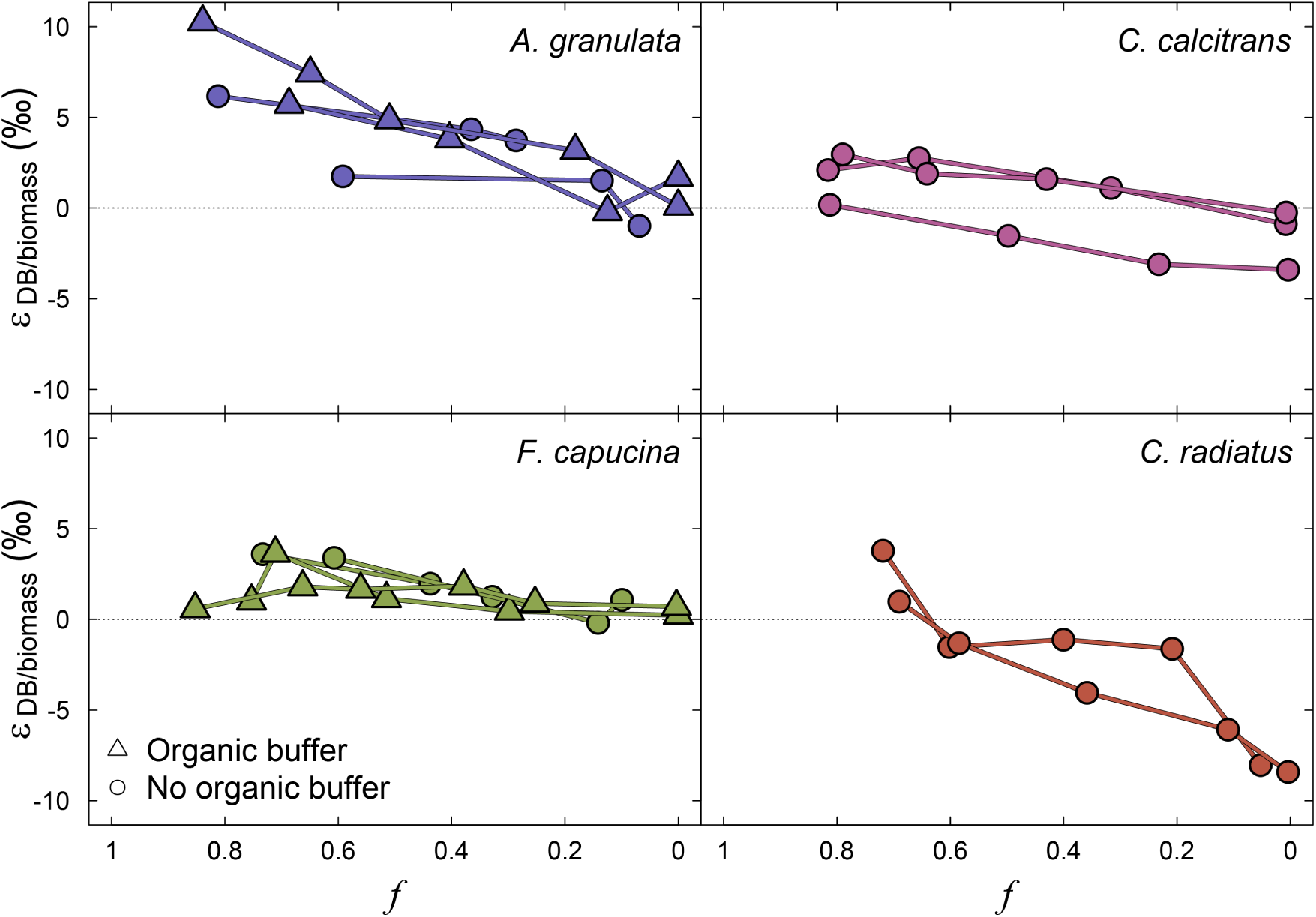
Isotopic offset between biomass and frustule-bound N (expressed as ε_DB/biomass_) progressing along the batch culture, shown as a function of the fraction of nitrate remaining (f). Points represent individual measurements, and lines connect sequential measurements from a single batch culture.

We further quantified the change in ε_DB/biomass_ along the *f* gradient, by calculating the slope of a linear regression for each batch culture, where the slope represents the rate of change of ε_DB/biomass_. When these slope values were plotted against the corresponding ε_biomass/NO3_ values (Fig. S4), a strong inverse relationship emerged. More precisely, cultures with less negative (i.e., higher) ε_biomass/NO3_ values exhibited a less pronounced slope, whereas those with more negative (i.e., lower) ε_biomass/NO3_ values displayed a stronger decrease in ε_DB/biomass_. This indicates that the extent to which frustule-bound N diverges from biomass δ^15^N is closely linked to the species-specific N isotope effect during nitrate assimilation.

C and N content (Table 1), and their ratio, could not explain variation in ε_DB/biomass_ within most species, except for *A. granulata*, where lower C contents and C/N ratios were associated with lower ε_DB/biomass_ values (p = 0.045 and p = 0.049, respectively).

### 3.5. Changes in biomass and frustule-bound δ^15^N values in response to high-δ^15^N nitrate additions

In an experiment to assess the incorporation of N isotope signatures from nitrate into biomass and frustule-bound N, we manipulated medium-nitrate δ^15^N values when roughly half of the initial nitrate pool had been consumed. Thereby the nitrate concentrations increased by only 1-2 µM, and spiked nitrate δ^15^N values upon ^15^N-labeled nitrate addition were between 85.5 ‰ and 94.4 ‰ (Fig. 6). Both biomass and diatom frustule-bound δ^15^N values clearly responded to the addition of ^15^N-labeled nitrate during batch cultures. Biomass δ^15^N values increased in close agreement with predictions for an accumulated product. Frustule-bound δ^15^N values also increased, but they remained consistently lower than those of biomass. The resulting ε_DB/biomass_ values were species-specific, with *A. granulata* showing smaller offsets compared to *F. capucina*. Upon complete nitrate consumption ε_DB/biomass_ values ranged from -6.4 to -8.0‰ for *A. granulata*, and from -23.6 to -23.8‰ for *F. capucina*. Using i) the δ^15^N values of samples collected before and shortly after addition of ^15^N-labeled nitrate, and ii) δ^15^N values after the spike addition, the relationship between biomass and frustule-bound δ^15^N values were assessed. Initially, the biomass-vs.-frustule-bound δ^15^N slopes were relatively shallow for both species, indicating that biomass δ^15^N values increased more strongly than frustule-bound δ^15^N values (Fig. 7). After these initial samples, the slope of this relationship approached 1 for *A. granulata* (0.93, p = 0.55 for slope ≠1), whereas for *F. capucina* the slope remained significantly lower than 1 (0.66, p = 0.04 for slope≠1). For *A. granulata*, the first after-spike samples were taken at roughly 3.6 and 5.5h after the ^15^N-nitrate addition, corresponding to a fractional nitrate consumption of 0.07*f* (Fig. S2). By contrast, for *F. capucina*, the first two samples were taken after 21.6 and 22.8h after spiking, corresponding to a fractional consumption of 0.19*f*. Throughout the experiment, empty frustule counts relative to frustules containing biomass were consistently low (< 3 %; Table S1).

**Figure 6:**
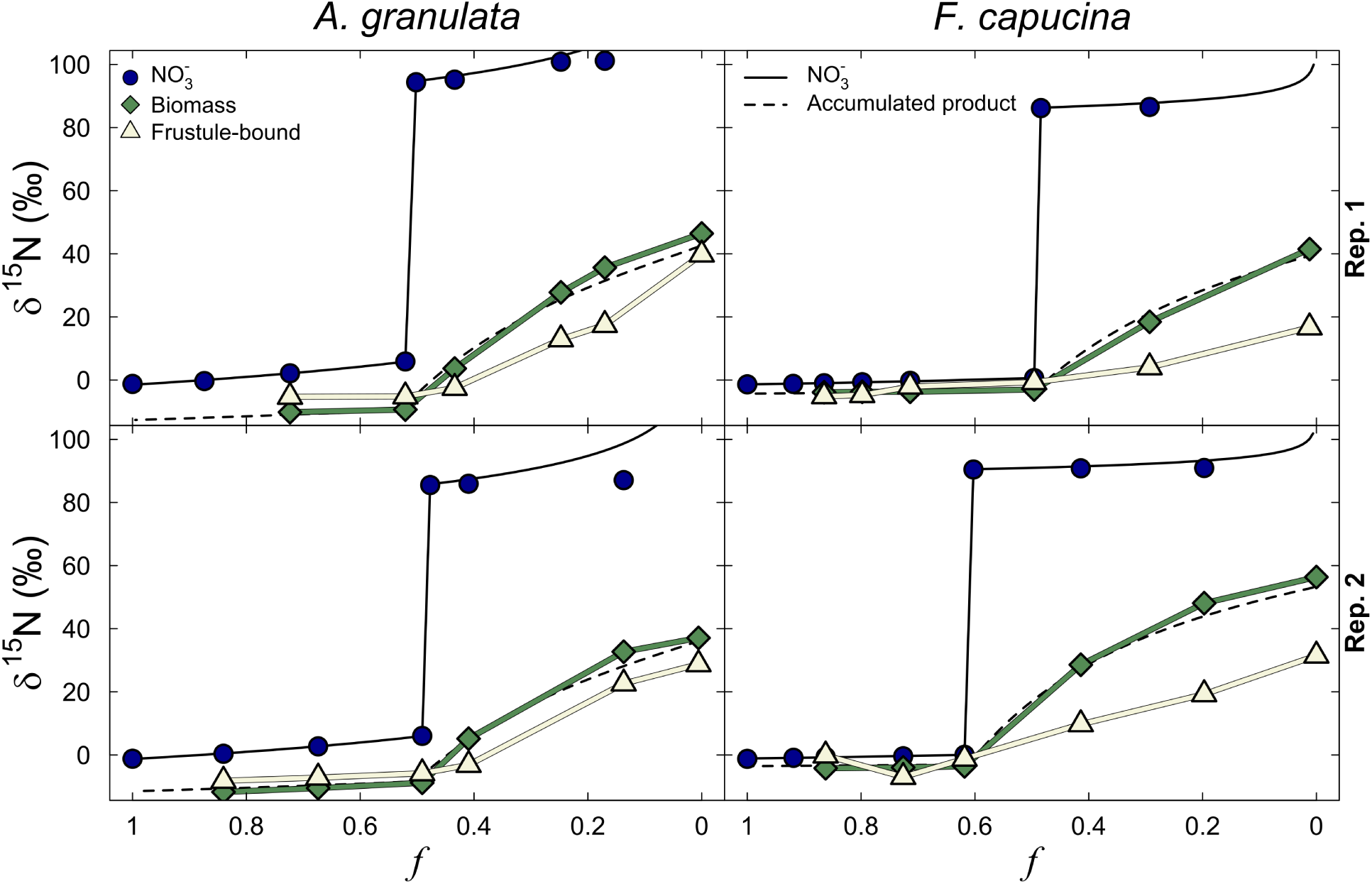
δ^15^N values of nitrate, biomass and frustule-bound N in response to the addition of high-δ^15^N nitrate (targeted nitrate δ^15^N 90‰) when roughly half of total nitrate had been consumed. Dashed lines represent estimated biomass δ^15^N values (accumulated product), calculated using an isotope mass balance approach following the addition of the nitrate δ^15^N spike.

**Figure 7:**
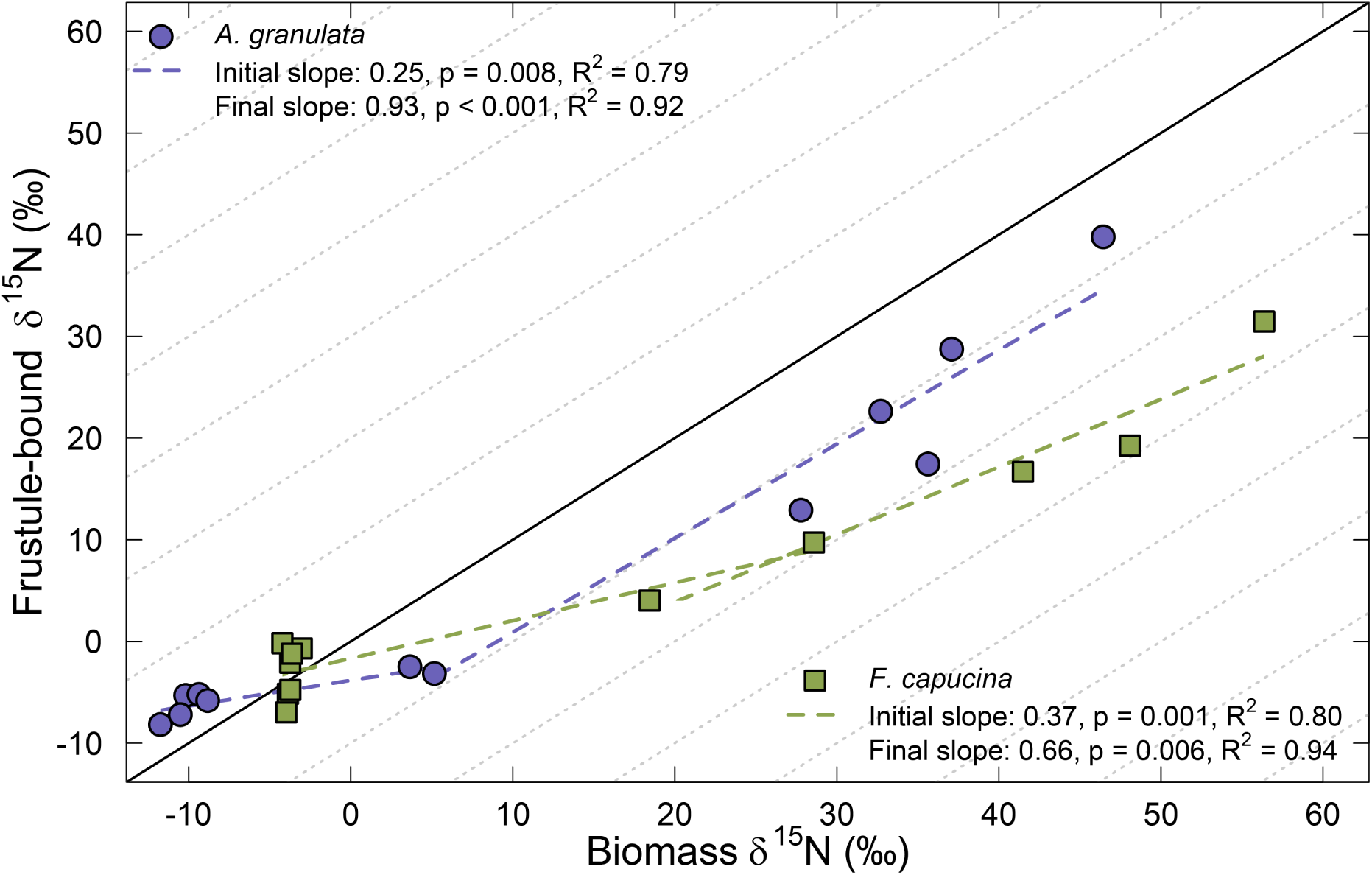
Frustule-bound δ^15^N values plotted against biomass δ^15^N values of samples taken in the experiment with ^15^N-labeled nitrate. Slopes (dashed lines) were determined on a species-specific basis and are distinguished by color. Initial slopes were calculated using the samples collected before the addition of ^15^N-labeled nitrate and the first two samples collected afterward. The ‘final slopes’ were calculated using only data from samples collected after the addition of ^15^N-labeled nitrate. The solid line represents a slope of 1 with an intercept 0. Dotted lines represent a slope of 1 with intercepts sequentially offset by 10 ‰.

## 4. Discussion

Meaningful interpretation of δ^15^N values of diatom frustule-bound organic N (δ^15^N_DB_) stored in sedimentary records relies on proper understanding of how N stable isotopes from the source nitrate pool are integrated into diatom biomass and subsequently into frustule-bound organic N. Here we present new data from mono-specific diatom cultures that illustrates the relationship among δ^15^N values of nitrate, biomass, and frustule-bound organic N in two freshwater and two marine diatom species. The results from these cultures show that δ^15^N_DB_ values can differ substantially from diatom biomass δ^15^N values, and that these differences depend on species as well as on the progression of the batch culture. Moreover, by experimentally manipulating nitrate δ^15^N values during batch culture incubations, we found that integration of the nitrate δ^15^N into biomass occurred more rapidly than into frustule-bound organic N, and the extent of this difference was species-dependent. However, these manipulations could only explain a small part of the discrepancies between biomass and frustule-bound δ^15^N values observed in the natural-abundance experiments, suggesting that the observed offsets are primarily driven by internal biochemical ^15^N fractionation.

### 4.1. Incorporation of nitrate δ^15^N signatures into biomass and frustule-bound N

Overall, we find that the ^15^N fractionation upon assimilation of nitrate (ε_biomass/NO3_) can vary substantially among species, but it is relatively stable within species under the conditions tested (Fig. 1), consistent with previous studies (Needoba et al., 2003). Moreover, measured biomass δ^15^N values agreed well with predicted biomass δ^15^N values calculated as accumulated product based on ε_biomass/NO3_ (Fig. 2 & 3). Thus, ^15^N fractionation of nitrate and subsequent incorporation into biomass followed predictable Rayleigh-type fractionation behavior in our batch cultures, as would be expected based on past research (Waser et al., 1998).

Previous studies using field-collected samples from the natural environment showed that, although biomass and frustule-bound δ^15^N values may be correlated with one another, neither was correlated with ambient nitrate δ^15^N values - likely due to substantial uptake of ammonium with distinct δ^15^N signatures (Morales et al., 2014; Robinson et al., 2020). In controlled batch cultures where nitrate is the only inorganic N source, both biomass δ^15^N and δ^15^N_DB_ would therefore be expected to follow the isotopic evolution of the nitrate pool, with biomass δ^15^N increasing according to ε_biomass/NO3_ and δ^15^N_DB_ anticipated to also track these changes. Indeed, across species and culture replicates, we find a significant relationship between δ^15^N values of frustule-bound N and those of biomass (Fig. 4). However, it is important to note that the slope of this relationship is significantly different from 1, and when the same comparison is made within species or within culture replicates, the relationship disappears completely. Moreover, the unexpectedly low explanatory power of this relationship (R^2^ = 0.32) indicates that only about one third of the observed variation in δ^15^N_DB_ values is explained by biomass δ^15^N values. Thus, the increase in biomass δ^15^N values induced by progressive nitrate consumption was not strictly mirrored by a corresponding increase in δ^15^N_DB_. This implies that δ^15^N_DB_ does not directly represent the δ^15^N of an accumulated product in batch cultures, and therefore it cannot be assumed to straightforwardly reflect biomass δ^15^N values.

### 4.2. The offset between frustule-bound and biomass δ^15^N changes with batch culture progression

We observed that the isotopic offset between frustule-bound and biomass δ^15^N values (ε_DB/biomass_) was not constant but decreased with batch culture progression (Fig. 5). This pattern primarily reflects the species-specific increase in biomass δ^15^N with fractional nitrate consumption, while δ^15^N_DB_ values remained comparatively stable (Fig. 3). Such relatively stable δ^15^N_DB_ values could imply that the organic N preserved in frustules originates from an N pool that remains unaltered throughout the batch culture. It has been proposed that the synthesis of frustule-bound organic N molecules can occur well before frustule formation, which may explain the low δ^15^N_DB_ values observed in *Cheatoceros* resting spores relative to frustules from vegetative cells (Dove et al., 2025). In batch culture experiments with natural-abundance nitrate δ^15^N levels, it is difficult to assess whether organic N in newly formed frustules undergoes variable biochemical ^15^N fractionation or isotopically reflects early culture stages, which is further discussed below. Irrespective of the underlying mechanism, the observed change in ε_DB/biomass_ provides useful context for interpreting batch culture results reported in previous studies.

Results from Horn et al. (2011) suggest a positive relationship between ε_DB/biomass_ and ε_biomass/NO3_. Similarly, our results also show that δ^15^N_DB_ values can be strongly offset from biomass δ^15^N values in species with large ε_biomass/NO3_ values, whereas the offsets are generally small in species with small ε_biomass/NO3_ values (Fig. 1 & 5). However, it is important to consider that ε_DB/biomass_ was not constant in our cultures (Fig. 3 & 5), and the magnitude of change in ε_DB/biomass_ during batch culture progression appears to be proportional to ε_biomass/NO3_ (Fig. S4). This suggests that processes driving variable ε_DB/biomass_ values could be related to physiological mechanisms involved in nitrogen assimilation and/or intracellular metabolism.

As silicification likely controls the frustule-bound organic compound matrix (Bridoux et al., 2012), and their synthesis timing (Kotzsch et al., 2017), silicification rates may affect incorporation of δ^15^N values into frustule-bound-N. The species used in our study represent a large range of silicification levels, with *C. radiatus* exhibiting Si:NO_3_ ^-^ uptake ratios greater than 4, compared to 0.4 to 0.7 for *C. calcitrans*, and 1.5 to 2.9 for the freshwater species (Table 1). In line with previous suggestions (Robinson et al., 2025), the heavily silicified *C. radiatus* was the only species whose δ^15^N_DB_ values were consistently lower than biomass δ^15^N values (Fig. 4 & 5). However, even though *A. granulata* and *F. capucina* had similar Si:NO_3_ ^-^ uptake ratios (Table 1), their observed ε_DB/biomass_ values differ substantially (Fig. 5). Moreover, ε_DB/biomass_ values were not consistent within single batch cultures, shifting from as high as +10 ‰ to around 0 ‰ in *A. granulata*, and from nearly +5 ‰ to -8 ‰ in *C. radiatus*. Depending on species, the silicification rates may change during batch culture progression as Si concentrations decrease (Kudo, 2003). However, we observed substantial shifts in ε_DB/biomass_ even under conditions where Si concentrations changed only modestly (Fig. 5; Fig. S1 & S2), and there was no clear relation with the replenishments of Si during the batch culture. Thus, our batch culture results do not support the proposed relationship between Si:NO_3_ ^-^ uptake ratios and ε suggested by Robinson et al. (2025).

In previous studies, it was difficult to gauge whether ε_DB/biomass_ values were stable or variable throughout batch cultures, since cultures were often terminated before complete nitrate consumption (Horn et al., 2011), or only a single sample was collected for δ^15^N_DB_ analyses (Jones et al., 2022; Robinson et al., 2025). Consequently, interpretations from previous culture studies may be confounded if ε_DB/biomass_ values cannot be assumed constant over the course of the batch culture. Importantly, our results imply that δ^15^N_DB_ values do not clearly follow the dynamics of an accumulating product in a Rayleigh-type system. The observed changes in ε_DB/biomass_ with batch culture progression are likely driven by (1) changes in ^15^N fractionation associated with shifts in biochemical reactions during N metabolism, and/or (2) differences in the timing of N isotope integration between δ^15^N_DB_ and biomass δ^15^N values through the use of intracellular N-reservoirs established early in the batch culture. To further explore the importance of the latter process, we conducted separate batch culture experiments in which ^15^N-labeled nitrate was added when roughly half of the initial nitrate remained, and traced its incorporation into biomass and frustule-bound N.

### 4.3. Slower nitrate isotope incorporation into the frustule-bound N pool does not explain εDB/biomass variation

The relatively constant δ^15^N_DB_ values observed during batch culture progression (Fig. 3) could suggest that ε_DB/biomass_ is mostly driven by the utilization of an isotopically stable N reservoir for the synthesis of frustule-bound organic N. To test this hypothesis, we conducted an experiment where nitrate δ^15^N values were manipulated during the batch culture. In this experiment, biomass δ^15^N values increased along the predicted Rayleigh path, whereas δ^15^N_DB_ values also increased but remained consistently lower than biomass δ^15^N values (Fig. 6; Fig. 7). One possible explanation for this offset could be a high or variable number of empty frustules relative to biomass-filled frustules; however, the abundance of empty frustules was consistently low (Table S1).

After addition of ^15^N-labeled nitrate, the distinctly lower δ^15^N values of frustule-bound N compared to biomass indicate that a portion of the organic N incorporated into frustules originates from before the δ^15^N manipulation. Moreover, the magnitude of this effect appears to be species-specific: frustule-bound δ^15^N values were substantially lower relative to biomass in *F. capucina* (−23.6 ‰) compared to *A. granulata* (−7.2 ‰) upon complete nitrate consumption. Additionally, the slope between biomass δ^15^N and δ^15^N_DB_ never reached 1 during the experiment time for *F. capucina*, whereas for *A. granulata* this slope approached 1 after only a few hours. Therefore, assimilation of ^15^N-labeled nitrate affects biomass δ^15^N values essentially instantaneously, while the pool of organic N used for frustule-bound organic N synthesis requires time to be replaced with newly assimilated, ^15^N-enriched N. An alternative explanation is that no new frustules were synthesized during the period when δ^15^N_DB_ values remained stable despite large change in biomass δ^15^N values; however, at least in *F. capucina* some new frustules must have formed during this time, as indicated by the observed Si consumption (Fig. S2). Thus, our results imply that frustule-bound organic N is derived from an intracellular N reservoir that integrates newly assimilated nitrate more slowly, leading to asynchronous incorporation of δ^15^N signals into frustule-bound N compared to bulk biomass.

Diatom-internal N reservoirs relevant to δ^15^N_DB_ values could take several forms: intracellular nitrate (Stief et al., 2022); monomers such as amino acids (Admiraal et al., 1986); N/C-storage compounds including urea and creatine-P (Allen et al., 2006; Parker et al., 2008); or the organic molecules (proteins and polyamines) involved in silicate precipitating prior to their incorporation into the frustule (Kotzsch et al., 2017; Tesson et al., 2017). Stored nitrate originating from early in the culture progression is unlikely to influence δ^15^N_DB_ values independently of biomass, since nitrate assimilation into organic compounds is not expected to occur via a temporally or spatially distinct pathway dedicated specifically to frustule-bound organic N synthesis.

Once assimilated into amino acids – the building blocks of proteins and polyamines, including those integrated into diatom frustules – N can be stored in various organic forms. In higher plants, amino acid turnover times can be less than one hour (Racusen and Foote, 1960), and in diatoms, substantial ^15^N enrichment in amino acids has been observed within one hour after ^15^ N-label addition (Smith et al., 2019). These rapid N turnover rates imply that amino acids are unlikely to retain isotopically distinct signatures over extended periods. Therefore, it is improbable that metabolically active amino acids constitute the source of delayed or asynchronous incorporation of ^15^N-labeled nitrate into frustule-bound organic N compared to biomass.

Many products of the diatom urea cycle can act as N-reservoirs, including urea and creatine-P (Allen et al., 2006; Parker et al., 2008). The consequences for frustule-bound δ^15^N values of these N-reservoirs depend on the extent to which such pools accumulate and are subsequently mobilized, processes that presumably vary with species identity and growth conditions. Assuming that utilization of these organic N-reservoirs occurs mainly under N-limiting conditions, it is unlikely that they were a major source of N for frustule-bound organic N synthesis, because N-limitation only developed toward the very end of the batch culture.

Pools of certain frustule-bound organic compounds may be established prior to frustule synthesis (Kotzsch et al., 2017; Tesson et al., 2017). If substantial amounts of these organic compounds are synthesized before frustule formation, a pool-size- and synthesis-rate-dependent amount of time would be required for them to be replaced with newly assimilated ^15^N-labeled N, thereby affecting the integration of nitrate δ^15^N values into the frustule-bound N pool.

Overall, the results from the ^15^N-labeled nitrate addition experiments (Fig. 6 & 7) imply that the organic N incorporated into frustules is derived from an organic N pool that is relatively quickly replaced by newly assimilated N. However, this time-window is clearly species-dependent, as slower-growing species required longer periods to replace these reservoirs with ^15^N-enriched N (Table 1, Fig. 7). Therefore, the incorporation of nitrate δ^15^N values into frustule-bound δ^15^N will lag behind their incorporation into biomass δ^15^N by a species-specific amount of time. This consideration is important for future batch culture experiments, as δ^15^N_DB_ values cannot be assumed to integrate over the exact same time window as biomass δ^15^N values.

With the slower integration of nitrate δ^15^N values into frustule-bound N compared to biomass, δ^15^N_DB_ values would still be expected to reflect the increase in biomass δ^15^N values in batch cultures – albeit with a temporal or *f*-related offset. Yet, in our batch cultures with natural-abundance nitrate δ^15^N values, δ^15^N_DB_ values showed hardly any increase at all (Fig. 3). Thus, to explain the observed changes in ε_DB/biomass_ values (Fig. 5), internal ^15^N fractionation must have varied with batch culture progression, which will be discussed next.

### 4.4. Potential biochemical ^15^N fractionation effects on frustule-bound δ^15^N values

Changes in growth conditions are an inherent feature of batch cultures, as nutrient concentrations decrease while cellular demand for nutrients (including inorganic C) increases with rising cell density. Increased cell density also causes self-shading, reducing light intensity in the culture. We found that biomass δ^13^C values progressively increased (or remained consistently elevated in the case of *C. radiatus*) (Fig. S3), which implies (1) a higher degree of inorganic C consumption (Degens et al., 1968; Deuser, 1970; Fry, 1996; Burkhardt et al., 1999), (2) induction of a carbon-concentration mechanism (Sharkey and Berry, 1985), and/or (3) enhanced C fixation via phosphoenolpyruvate carboxylase in response to increased ammonium assimilation (Guy et al., 1989). Thus, the observed biomass δ^13^C patterns illustrate that the progressive changes in growth conditions that inherently occur during batch culture progression affect diatom physiology. Overall, the initially higher, and subsequently decreasing, frustule-bound δ^15^N values relative to biomass (Fig. 5) could be a result of growth-condition-driven changes in (1) the frustule-bound organic compound matrix, (2) ^15^N-fractionating reactions during frustule-bound organic compound synthesis, and/or (3) ^15^N-fractionating reactions within amino acid synthesis pathways. In the following, we address potential mechanisms by which ε_DB/biomass_ may – or may not – have changed.

Regarding (1): If the frustule-bound organic compound matrix changed during the batch cultures, associated with a shift from N-rich precursors with high δ^15^N values to precursors with lower δ^15^N values, this would be reflected in δ^15^N_DB_. However, it is currently unknown whether the composition of the frustule-bound organic matrix is plastically regulated within species (i.e., whether it varies within species in response to growth conditions). Yet, because the frustule-bound polyamine composition is highly species-specific (Bridoux and Ingalls, 2010; Bridoux et al., 2012), it is more likely that the frustule-bound organic compound matrix is governed primarily by diatom phylogeny rather than by growth conditions.

Regarding (2): For proteins, ribosomal biosynthesis from amino acids is generally assumed not to introduce isotopic discrimination (Werner and Schmidt, 2002). However, in higher plants, polyamine δ^15^N values have been found to be substantially higher than those of their amino acid precursors, suggesting considerable potential for ^15^N fractionation during polyamine biosynthesis (Yoneyama et al., 1998). Moreover, in diatoms, methylation of amine groups in frustule-bound silaffins and polyamines has also been documented (Sumper and Brunner, 2008; Kröger and Poulsen, 2008; Bridoux and Ingalls, 2010), and since such reactions involve the breaking of N-H and forming of N-C bonds, they may introduce ^15^N fractionation. This isotopic fractionation could be expressed if methylation occurs at branching points within the synthesis pathway, and/or if the substrate-product reaction is incomplete. Increased methylation has been shown to enhance the ability of polyamines to precipitate silicic acid (Robinson et al., 2007), and the degree of methylation varies among species (Bridoux et al., 2012). However, whether changes in the degree of methylation of frustule-bound organic-N can occur *within* a species, thereby producing variable ^15^N fractionation and contributing to the observed changes in ε_DB/biomass_, remains unclear.

Regarding (3): To identify potential ^15^N fractionation processes prior to the synthesis of frustule-bound organic compounds, it is important to understand how these molecules are synthesized, and from which precursors. Polyamines are synthesized from a putrescine or ornithine backbone and from polypropylene-imine (PPI) chains of species-dependent lengths, which are ultimately derived from methionine synthesized from aspartate (Bridoux and Ingalls, 2010; Lin et al., 2024). Thus, for polyamines composed of more than two PPI units, methionine represents the dominant source of N. By comparison, frustule-bound proteins such as silaffins and silacidins are rich in serine (Poulsen et al., 2003; Richthammer et al., 2011), which is produced primarily during photorespiration, whereas other frustule-associated proteins have been shown to be enriched in glutamine and asparagine (Kotzsch et al., 2017), amino acids more closely related to N assimilation. Assuming that metabolic fluxes influence the extent of isotopic fractionation (Tcherkez, 2010), diatom metabolism likely plays an important role in shaping amino acid δ^15^N values (Fig. 8) and, consequently, the δ^15^N of frustule-bound organic N compounds. Understanding the variation among amino acid δ^15^N values and the metabolic processes influencing them may therefore provide a comprehensive framework for identifying potential biochemical controls on frustule-bound δ^15^N.

**Figure 8:**
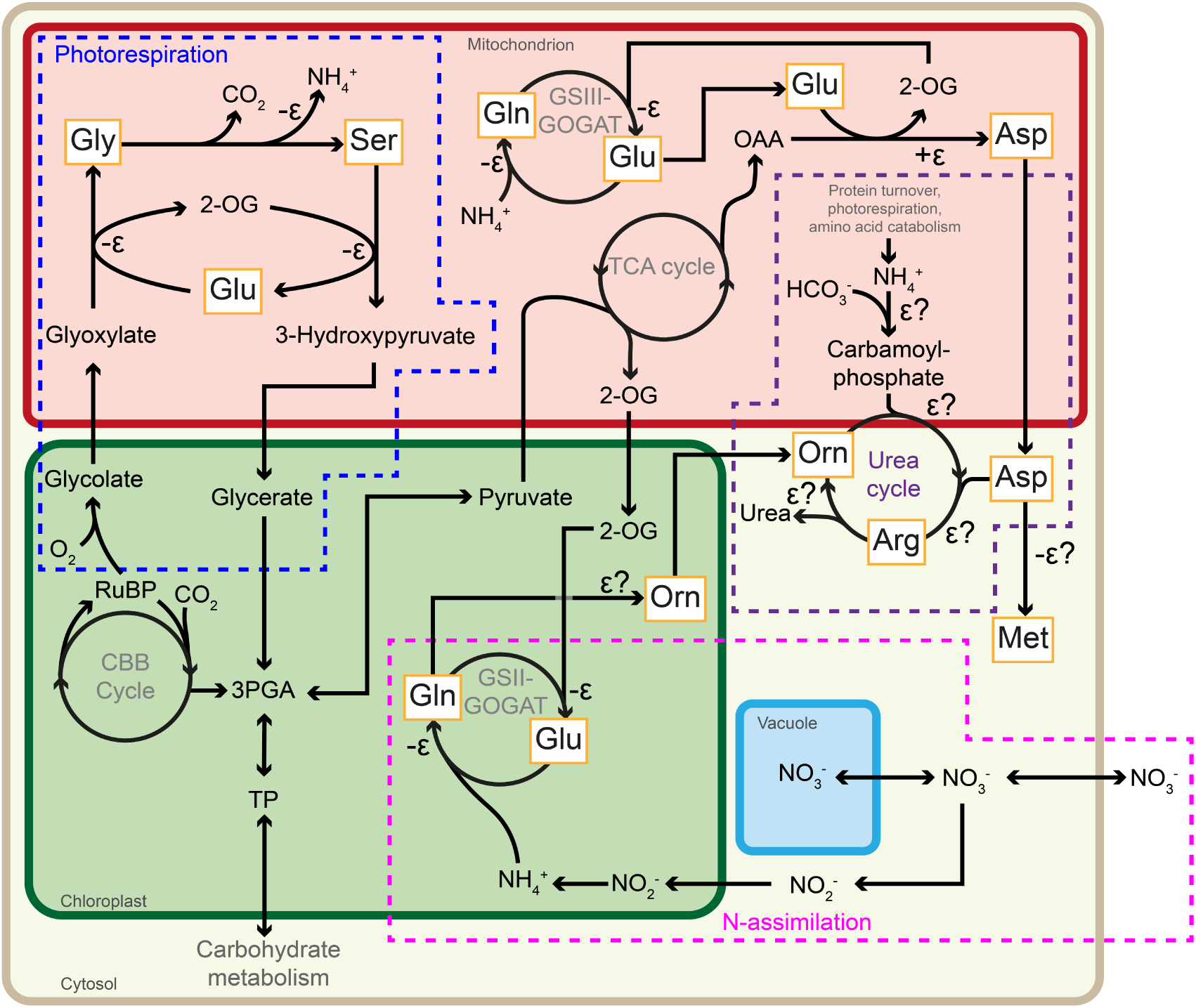
Schematic, simplified overview of the interaction between diatom C and N metabolism. The main areas of interaction are N-assimilation (pink), photorespiration (blue), and the urea cycle (purple). Reactions involving N-containing organic compounds are annotated with ε and a sign prefix, where -ε indicates a reaction resulting in lower product δ^15^N values, and +ε indicates the opposite. In cases where the ^15^N fractionation is unknown, ‘ε?’ is shown. In the figure, distinctions are made between GS-GOGAT located in the plastid (GSII-GOGAT) involved in N-assimilation and GS-GOGAT located in the mitochondrion (GSIII-GOGAT) involved in ammonium recycling. Abbreviations: 2-OG, 2-oxoglutarate; 3PGA, 3-phosphoglycerate; Arg, arginine; Asp, aspartate; CBB cycle, Calvin-Benson-Bassham cycle; Gln, glutamine; Glu, glutamate; GOGAT, glutamate synthase; GS, glutamine synthetase; Gly, glycine; Met, methionine; Orn, ornithine; OAA, oxaloacetate; RuBP, ribulose bisphosphate; TCA cycle, tricarboxylic acid cycle; TP, triose-phosphate. Information compiled from multiple publications (Werner & Schmidt, 2002; Kroth et al., 2008; Tcherkez & Hodges, 2008; Smith et al., 2019).

Transamination and deamination reactions in the photorespiration cycle lead to relatively low δ^15^N values of serine in photosynthetic organisms (Fig. 8) (Tcherkez and Hodges, 2008; Gauthier et al., 2013; McCarthy et al., 2013; McMahon and McCarthy, 2016; Takizawa et al., 2017). Although photorespiration strongly impacts the δ^15^N values of amino acids relevant for frustule-bound protein synthesis (Fig. 8) (Tcherkez and Hodges, 2008; Tcherkez, 2010), variation in photorespiratory flux in diatoms is likely minimized by carbon-concentrating mechanisms (Clement et al., 2016) and the increased specificity of the RubisCO enzyme for CO_2_ compared to O_2_ (i.e., which reduces oxygenation reactions and thereby suppresses photorespiration; Tortell, 2000; Kroth et al., 2008). However, photorespiration in diatoms can be induced under high-light conditions (Parker et al., 2004), indicating that variable photorespiratory flux is possible, and could therefore affect the δ^15^N values of amino acids used in the synthesis of frustule-bound organic compounds.

Methionine δ^15^N values in photosynthetic organisms are typically substantially lower than those of glutamate (Ishikawa et al., 2018). Considering that aspartate δ^15^N values are generally higher than those of glutamate (Tcherkez and Hodges, 2008), the relatively low methionine δ^15^N values imply strong ^15^N fractionation during its synthesis or utilization. Diatoms possess a cobalamin (vitamin B_12_)-dependent methionine synthase enzyme, MetH, and certain species also contain a cobalamin independent form, MetE (Bertrand et al., 2012). The latter is considerably less efficient than the cobalamin-dependent enzyme. Thus, depending on whether MetH or MetE is used, substantial ^15^N fractionation may occur during methionine synthesis and processing, reflecting vitamin B_12_ availability in the medium. Our batch cultures contained cobalamin concentrations substantially higher than those in the F/2 medium used in previous studies (Horn et al., 2011; Robinson et al., 2025), making it unlikely that vitamin B_12_ levels were low enough to influence methionine δ^15^N values. Nevertheless, the oceanographic relevance of vitamin B12 and its potential effect on methionine synthesis should be carefully considered in δ^15^N_DB_ related research.

An important source of N in the urea cycle is carbamoyl phosphate, formed from bicarbonate and ammonium (Fig. 8). Transcripts of the carbamoyl phosphate synthase gene (unCPS) are more abundant under high-light compared to low-light conditions (Bender et al., 2012), indicating sensitivity of N metabolism to light. Moreover, unCPS transcript abundance shown in Bender et al. (2012) correlates with transcripts for glutamine synthesis in both the chloroplast and mitochondrion, linking N-assimilation to the urea cycle (Fig. 8). Unlike higher plants, in which ammonium released during photorespiration and amino acid catabolism is predominantly re-assimilated in the chloroplast, in diatoms this ammonium can also be re-assimilated via carbamoyl phosphate synthesis and subsequently processed through the urea cycle (Allen et al., 2006, 2011; Parker et al., 2008). Assuming that deamination reactions releasing ammonium impart isotopic fractionation (Tcherkez and Hodges, 2008; Takizawa et al., 2017), this pathway could shuttle N with metabolism-imparted distinct δ^15^N signatures into the urea-cycle products, which can then be redistributed to other amino acids.

Another notable result is the observed relationship between ε_biomass/NO3_ values and the temporal change in ε_DB/biomass_ over the course of batch culture progression (Fig. S4). During nitrate assimilation, nitrate is converted to nitrite and subsequently to ammonium. The conversion of nitrate to nitrite by nitrate reductase (NR) is typically the limiting step in this sequence. The magnitude of the associated ^15^N fractionation, expressed as ε_biomass/NO3_, depends on the balance between nitrate influx and efflux across the cell membrane (Granger et al., 2010; Chen et al., 2024). Because ε_biomass/NO3_ is largely governed by transport/diffusion and reduction kinetics, it is unlikely to directly cause the variation observed in ε_DB/biomass_. Instead, the variable relationship between the two likely reflects species-specific differences in N metabolism and the degree of internal N-isotope fractionation during biosynthetic processing of N. This interpretation is supported by the relation between amino acid δ^15^N values and N-assimilation rates in higher plants (Tcherkez, 2010), and the connection between N-assimilation pathways and the urea cycle in diatoms (Bender et al., 2012).

Ultimately, based on our experiments alone, the specific mechanism underlying changes in ε_DB/biomass_ could not be determined. Nevertheless, our results strongly suggest that frustule-bound δ^15^N values are sensitive to changes in growth conditions (such as light intensity and inorganic C availability) that influence metabolically driven ^15^N fractionation during intracellular processing of organic N. We thus hypothesize that alterations in metabolic fluxes within the N-assimilation, photorespiration, and urea-cycle pathways may play particularly important roles in determining ^15^N fractionation of amino acids used for frustule-bound organic compound synthesis.

### 4.5. Implications for the diatom frustule-bound nitrogen isotope paleo-proxy

The observed slower integration of nitrate δ^15^N values into frustule-bound organic N compared to bulk biomass is likely not important for paleorecords, because the timescale of this lag - on the order of a few hours to at most a few days - is negligible relative to the multiple-year to millennial integration represented in sedimentary δ^15^N_DB_. More importantly, for *A. granulata*, we showed that after this brief lag period, δ^15^N_DB_ values follow those of biomass relatively closely (Fig. 7), confirming that δ^15^N_DB_ values can, in principle, faithfully track changes in nitrate δ^15^N across a wide range of values.

Although this short-term time lag is negligible for paleo-applications, our results also show that δ^15^N_DB_ may diverge from biomass δ^15^N for reasons other than timing alone. Historically, δ^15^N_DB_ values have been interpreted as direct reflections of biomass δ^15^N values. However, the present work reveals substantial ambiguity in how closely δ^15^N_DB_ values track biomass δ^15^N, consistent with observations from other diatom culture studies (Horn et al., 2011; Jones et al., 2022; Robinson et al., 2025). We hypothesize that species-specific and growth-condition-dependent interactions between diatom C and N metabolism and the associated ^15^N fractionation can lead to variability in δ^15^N values of frustule-bound organic N that is not directly predictable from biomass δ^15^N values alone. Whether such interactions also impact frustule-bound δ^15^N values under natural environmental conditions remains to be determined.

In sedimentary records from the Southern Ocean, diatom frustule-bound δ^15^N has been shown to vary on glacial/interglacial timescales, with δ^15^N_DB_ values up to 4 ‰ higher during glacial periods compared to interglacials (Robinson et al., 2004; Studer et al., 2015). Glacial/interglacial cycles are characterized by large changes in atmospheric CO_2_ concentrations and marine biological productivity, with both being lower in the Antarctic Ocean during glacial periods (Mortlock et al., 1991; Petit et al., 1999; Studer et al., 2015). The elevated glacial δ^15^N_DB_ values have generally been attributed to reduced nitrate supply, leading to increased fractional nitrate consumption and thus higher δ^15^N values incorporated into frustule-bound organic N (Robinson et al., 2004; Studer et al., 2015). In contrast to the natural environment, the batch culture experiments in our study were conducted under near-unlimited nutrient conditions until nitrate concentrations approached complete consumption. Despite this, ε_DB/biomass_ was highly variable, suggesting that ε_DB/biomass_ was not directly driven by external nutrient status, but rather by other factors, such as light conditions, inorganic C supply, and/or cell density. A previous study highlighted that the isotope effect of nitrate assimilation in the Antarctic Ocean is insensitive to mixed-layer depth and thus, light availability (Fripiat et al., 2019). Moreover, improved light conditions are unlikely to contribute to the observed glacial increase in frustule-bound δ^15^N in paleorecords, because light-driven increases in nitrate consumption would have been associated with higher productivity, opposite to what is observed in the Antarctic Ocean during glacial periods (Studer et al., 2015; Ai et al., 2020).

With respect to atmospheric CO_2_, it is important to consider whether diatom growth could have been C-limited during glacial periods, potentially inducing changes in diatom metabolic pathways relevant to internal biochemical ^15^N fractionation. Although phytoplankton growth is often assumed not to be limited by inorganic C in the natural environment, diatoms can experience CO_2_ limitation during dense blooms, especially in air-equilibrated surface ocean waters during glacial periods when atmospheric CO_2_ concentrations were ∼200 ppm (Riebesell et al., 1993). Since our results suggest that the generation of frustule-bound δ^15^N signals can be sensitive to growth conditions, this aspect should be carefully considered when interpreting sedimentary δ^15^N_DB_ records.

The continued development of the δ^15^N_DB_ proxy should include systematic characterization of how diatom metabolic processes affect frustule-bound δ^15^N values. As discussed above, compound-specific ^15^N-isotope analysis of biomass-derived amino acids will provide vital information linking diatom metabolism to the isotopic fractionation mechanisms that shape δ^15^N_DB_. Additionally, δ^15^N measurements of amino acids derived from frustule-bound organic N could be highly informative for distinguishing between shifts in source-N δ^15^N values (where all amino acid δ^15^N values change similarly) and metabolically driven biochemical ^15^N fractionation effects, where amino acid δ^15^N values change relative to one another. Such approaches have already been applied to mollusk shells from sediment archives (Vokhshoori et al., 2023), and extending them to diatom frustules would provide significant new insights for refining the δ^15^N_DB_ paleo-proxy.

## 5. Conclusion

The results presented here demonstrate that diatom biomass and frustule-bound δ^15^N values can become decoupled in batch cultures, and that this decoupling cannot be explained by differences in isotopic integration timing. We show that δ^15^N_DB_ is indeed driven by δ^15^N values of assimilated nitrate in ^15^N-labeled nitrate experiments; however, under natural-abundance conditions, ^15^N fractionation during intracellular processing of organic N substantially influences δ^15^N_DB_. We hypothesize that this offset arises from growth-condition-dependent shifts in metabolic fluxes, which can influence ^15^N fractionation during amino acid synthesis and the formation of frustule-bound organic compounds. Future ground-truthing efforts should therefore aim to quantify the extent and drivers of decoupling between bulk biomass and frustule-bound δ^15^N under controlled varying environmental conditions. Such experiments would improve our understanding of how δ^15^N_DB_ values respond to physiological variability during diatom growth, ultimately enhancing the reliability of the δ^15^N_DB_ paleo-proxy for reconstructing N cycling and environmental change in aquatic systems over long time scales.

## Author contributions

**Jochem Baan**: Conceptualization, Data curation, Investigation, Formal analysis, Methodology, Visualization, Writing – original draft, Writing – review and editing; **Moritz F. Lehmann**: Conceptualization, Resources, Supervision, Writing – review and editing; **Anja S. Studer**: Conceptualization, Funding acquisition, Investigation, Methodology, Project administration, Supervision, Writing – review and editing.

## Data availability

The resulting dataset from this study is available on the *Zenodo* repository (https://doi.org/10.5281/zenodo.16994169)

## Declaration of competing interest

The authors declare that they have no known competing financial interests or personal relationships that could have appeared to influence the work reported in this paper.

## Acknowledgements

Funding for this study was provided by the Swiss National Science Foundation (Grant 200021_200766 to A.S.S.) and by resources from the University of Basel. We would like to thank Marta Reyes (Eawag) for providing freshwater diatom cultures and help with diatom culturing techniques; Thomas Kuhn (University of Basel) for help with Flash-EA IRMS analyses of biomass samples and general laboratory support; Klaus Schläppi (University of Basel) for providing access to SANYO climate chambers; Ansgar Kahmen (University of Basel), Günter Hoch (University of Basel), and Georges Grün (University of Basel) for access to, and setting up phytotron growth chambers; Gregory de Souza (ETH Zürich) for help with silicate concentration analysis; and Guillaume Tcherkez (University of Angers), Roland Werner (ETH Zürich) and Peter Kroth (University of Konstanz) for helpful discussions of biochemical ^15^N fractionation and diatom metabolism.

## Appendix A. Supplementary material

**Supplementary Table S1:** Percentage of empty frustules for each sample collected during batch cultures spiked with ^15^N-labeled nitrate.

**Supplementary Figure S1:** Silicate concentration, fraction of nitrate remaining in the medium (*f*) and pH over time during batch culture progression.

**Supplementary Figure S2:** Silicate concentration, fraction of nitrate remaining in the medium (*f*) and pH over time during batch culture progression, where ^15^N-labeled nitrate was added after approximately half of the initial nitrate pool had been consumed.

**Supplementary Figure S3:** Biomass δ^13^C values plotted against *f*.

**Supplementary Figure S4:** Slopes of the linear regression between ε_DB/biomass_ and fraction of nitrate remaining in the medium (*f*) plotted against ε_biomass/NO3_ values.

## Notes

### Competing Interest Statement

The authors have declared no competing interest.

### Summary of Updates

General textual revisions throughout the manuscript. Removal of author that was no longer interested in co-authorship.

