## Supplemental Table S1 and Figures S1-4 for "Batch culture and species effects modulate the decoupling between diatom frustule-bound and biomass nitrogen isotope signatures"

Supplementary Table S1 and Figures S1-4

*Table S1: Percentage of empty frustules for each sample collected during batch cultures spiked with $^{15}$N-labeled nitrate. Values are expressed as the mean percentage ±sd, based on three to four replicate counts of empty frustules out of 100 frustules.*

| Species | Batch culture replicate | Time after inoculation (d) | % empty frustules (mean ±sd) | Pre-/post- $^{15}$N-labeled nitrate spike |
|---|---|---|---|---|
| *A. granulata* | 5 | 11.8 | 2.3 ±0.5 | Pre- |
| | | 12.8 | 2.0 ±0.8 | Pre- |
| | | 13.1 | 2.0 ±1.4 | Post- |
| | | 13.8 | 1.0 ±0.8 | Post- |
| | | 14.0 | 0.7 ±0.5 | Post- |
| | | 14.8 | 1.0 ±0.0 | Post- |
| | 6 | 8.8 | 0.3 ±0.5 | Pre- |
| | | 10.0 | 0.7 ±0.5 | Pre- |
| | | 10.8 | 0.7 ±0.9 | Pre- |
| | | 11.0 | 0.3 ±0.5 | Post- |
| | | 11.8 | 0.3 ±0.5 | Post- |
| | | 12.0 | 0.7 ±0.5 | Post- |
| *F. capucina* | 5 | 15.8 | 1.3 ±0.5 | Pre- |
| | | 17.0 | 0.7 ±0.5 | Pre- |
| | | 17.9 | 0.3 ±0.5 | Post- |
| | | 19.0 | 1.3 ±1.2 | Post- |
| | | 19.9 | 1.0 ±0.8 | Post- |
| | | 21.0 | 0.5 ±0.5 | Post- |
| | 6 | 15.8 | 1.3 ±0.5 | Pre- |
| | | 17.0 | 0.3 ±0.5 | Pre- |
| | | 17.9 | 0.7 ±0.5 | Pre- |
| | | 19.0 | 1.3 ±0.9 | Post- |
| | | 19.9 | 1.7 ±0.5 | Post- |
| | | 21.0 | 0.3 ±0.5 | Post- |

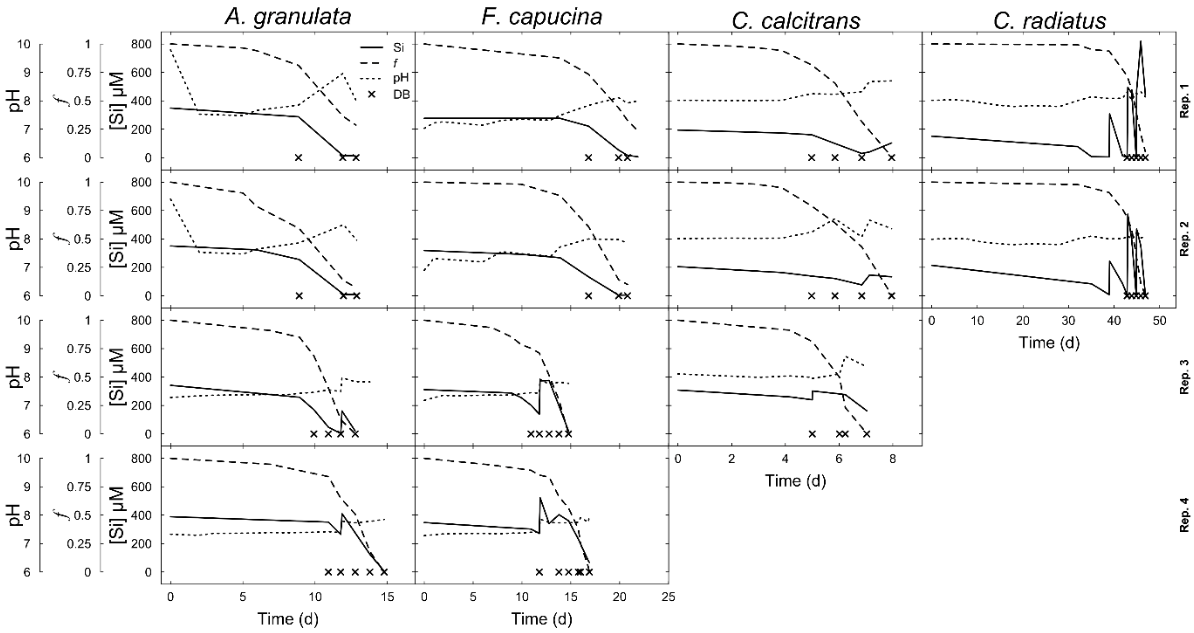


*Figure S1: Silicate concentration, fraction of nitrate remaining in the medium (f) and pH over time during batch culture progression. Time points at which biomass was harvested for analyses are indicated by ‘x’.*

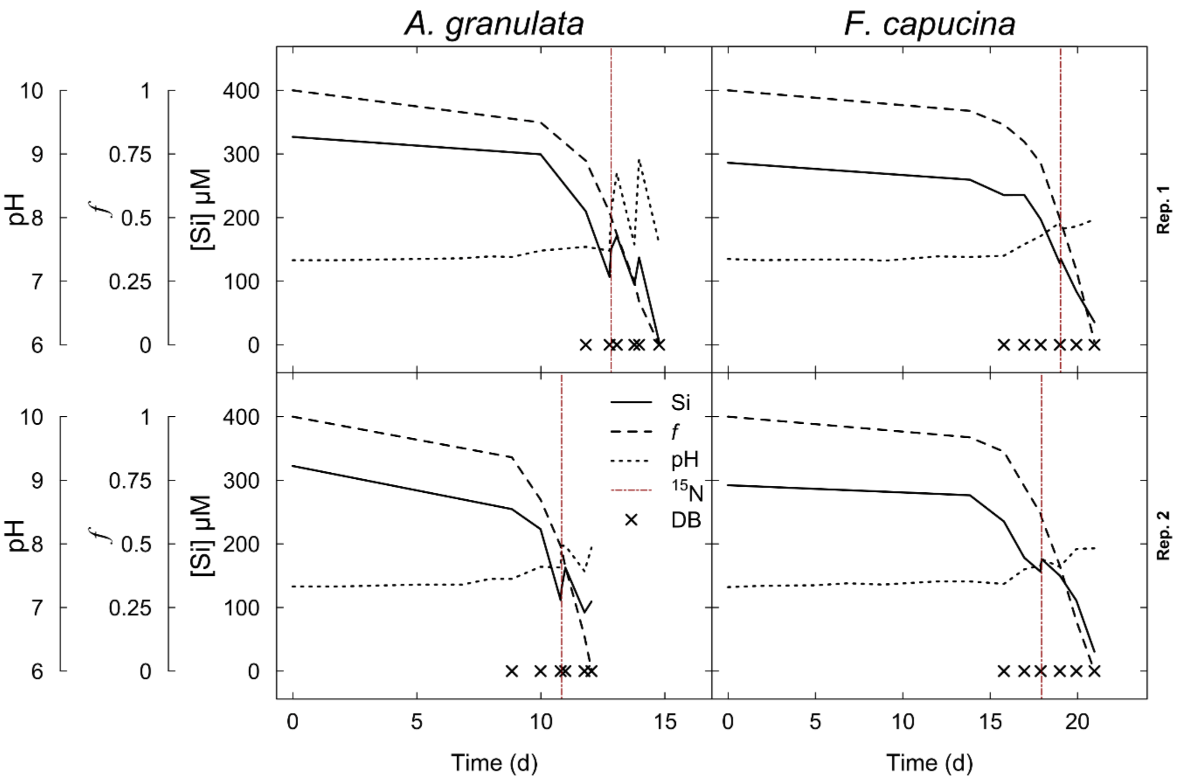


*Figure S2: Silicate concentration, fraction of nitrate remaining in the medium (f) and pH over time during batch culture progression, where $^{15}N$-labeled nitrate was added after approximately half of the initial nitrate pool had been consumed. Time points at which biomass was harvested for analyses are indicated by 'x'. The time point at which $^{15}N$-labeled nitrate was added is indicated by a vertical red line.*

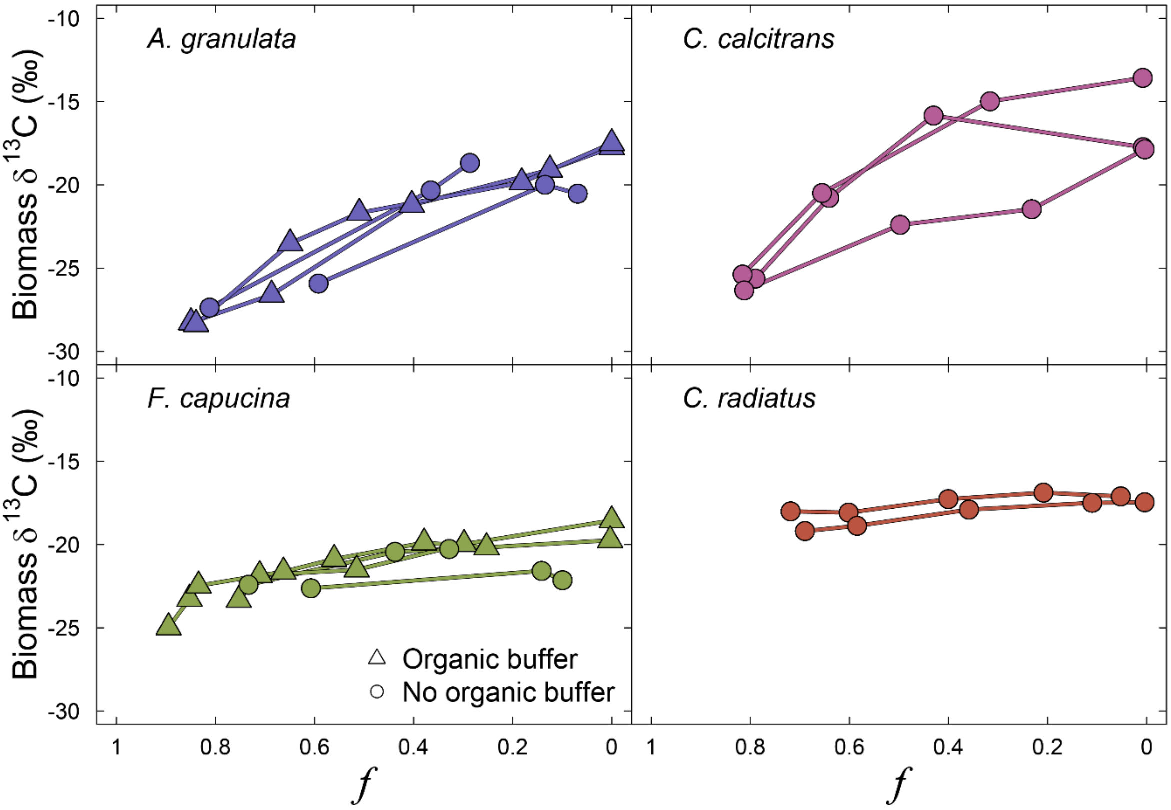


*Figure S3: Biomass $\delta^{13}C$ values plotted against f as an indicator for batch culture progression. Lines connect sequential samples from the same batch culture.*

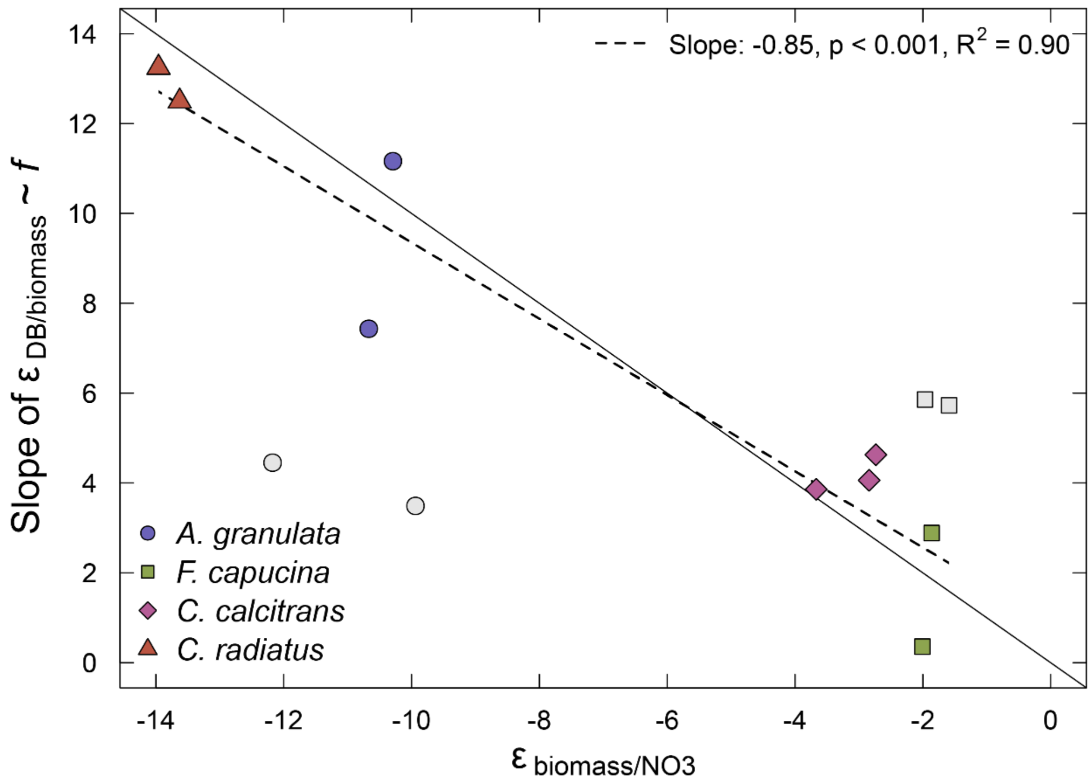


*Figure S4: Slopes of the linear regression between $\varepsilon_{DB/biomass}$ and fraction of nitrate remaining in the medium (f) plotted against $\varepsilon_{biomass/NO3}$ values. The dashed line is the linear regression while the solid line represents a slope of -1. Light gray points are from batch cultures with less than four individual frustule samples and are not used in this linear regression analysis.*
